# The fungal root endophyte *Serendipita vermifera* displays inter-kingdom synergistic beneficial effects with the microbiota in *Arabidopsis thaliana* and barley

**DOI:** 10.1101/2021.03.18.435831

**Authors:** Lisa K. Mahdi, Shingo Miyauchi, Charles Uhlmann, Ruben Garrido-Oter, Gregor Langen, Stephan Wawra, Yulong Niu, Senga Robertson-Albertyn, Davide Bulgarelli, Jane E. Parker, Alga Zuccaro

## Abstract

Plant root-associated bacteria can confer protection against pathogen infection. By contrast, the beneficial effects of root endophytic fungi and their synergistic interactions with bacteria remain poorly defined. We demonstrate that the combined action of a fungal root endophyte from a widespread taxon with core bacterial microbiota members provides synergistic protection against an aggressive soil-borne pathogen in *Arabidopsis thaliana* and barley. We additionally show early inter-kingdom growth promotion benefits which are host and microbiota composition dependent.

**Highlights:**

- The root endophytic fungus *Serendipita vermifera* can functionally replace core bacterial microbiota members in mitigating pathogen infection and disease symptoms.
- *S. vermifera* additionally stabilizes and potentiates the protective activities of root-associated bacteria and mitigates the negative effects of a non-native bacterial community in *A. thaliana*.
- Inter-kingdom synergistic beneficial effects do not require extensive host transcriptional reprogramming nor high levels of *S. vermifera* colonisation.
- Inter-kingdom protective benefits are largely independent of the host while synergism leading to early inter-kingdom growth promotion is driven by host species and microbiota composition.

## Introduction

Plant pathogenic fungi limit crop productivity globally. These threats are expected to increase with global warming (Delgado-Baquerizo et al., 2020). Decades of advances in agrochemicals and plant breeding have expanded farmers’ toolkits with fungicides and resistant varieties to limit detrimental effects of these organisms on crop yield. Yet, current tools are becoming environmentally unsustainable or ineffective against rapidly evolving pathogens (Delgado-Baquerizo et al., 2020). A key example of this scenario is represented by the soil-borne plant pathogen *Bipolaris sorokiniana* (syn. *Cochliobolus sativus*, hereafter *Bs*), the causal agent of spot blotch and common root rot diseases that threaten cereal production in warm regions (Delgado-Baquerizo et al., 2020; Duveiller and Gilchrist, 1994; Manamgoda et al., 2014). Root rot normally originates from inoculum carried on the seed or from soil-borne conidia, but the fungus can infect plants at any developmental stage. However, as the importance of root-inhabiting pathogenic fungi has often been underestimated, very little is known about the molecular mechanism behind the detrimental interaction of *Bs* with roots (Sarkar et al., 2019).

Microbial communities living at the root-soil interface, collectively referred to as the plant root microbiota, have gained center-stage in pathogen protection (Bulgarelli et al., 2015). Past studies across a variety of plant species employed environmental sampling or controlled conditions in the field and laboratory to characterise the root microbiota (Duran et al., 2018; Edwards et al., 2015; Fitzpatrick et al., 2018; Lundberg et al., 2012; Thiergart et al., 2020a), with an overall greater focus on bacteria than on filamentous fungi (Whipps, 2001). Microbial diversity and abundance gradually decrease between the soil and vicinity of the root (rhizosphere), and further between the rhizosphere and root internal compartments (endosphere). Moreover, a number of bacterial taxa (e.g., Proteobacteria, Actinobacteria, Bacteroidetes and Firmicutes) consistently occur in the root endosphere of different examined plant species (Fitzpatrick et al., 2018). This latter feature underpins the “bacterial core microbiota” concept, in which strains from specific taxa are commonly selected as endophytes across plant species, soil types and environmental conditions (Lemanceau et al., 2017). By contrast, studies of geographically distinct populations of *Arabis alpina* and *Arabidopsis thaliana* (hereafter Arabidopsis) showed that few fungal taxa are prevalent in the root endosphere, and that endophytic fungal communities are strongly influenced by location and climate (Almario et al., 2017; Thiergart et al., 2020a).

The functions and benefits of root microbiota members in the context of abiotic or biotic stresses have been extensively investigated under laboratory conditions using single microbial strains and, more recently, synthetic bacterial communities (SynComs) (Vorholt et al., 2017). Several bacterial and fungal isolates have the capacity to directly increase plant biomass via growth hormone production and/or by providing plants with limiting macro- or micro-nutrients (Almario et al., 2017; Franken, 2012; Harbort et al., 2020; Hermosa et al., 2012; Hiruma et al., 2016; Spaepen et al., 2007). Although diseases caused by pathogens have been shown to be directly or indirectly reduced by the addition of single or multiple beneficial microbes (Pieterse et al., 2014; Vlot et al., 2020), how fungal root microbiota members with beneficial functions influence and are influenced by bacterial colonisation remains less understood.

Sebacinales fungi (Basidiomycetes) are a remarkable group of plant mutualists with worldwide occurrence in soils and as endophytes. While single Sebacinales strains can interact with roots in the absence of differentiated structures, they can also form specialized interactions with distinctive morphological characteristics on relevant hosts, as in orchid- or ectomycorrhiza symbioses (Weiss et al., 2016a). Root colonisation by these fungi improved host growth and development, increased grain yield and enhanced root phosphate uptake in several plant species (Fesel and Zuccaro, 2016; Oberwinkler et al., 2013; Zuccaro, 2020; Zuccaro et al., 2014). The positive effects of Sebacinales on the host plant extend well beyond growth and development and cannot be explained by enhanced host nutrition alone (Oberwinkler et al., 2013; Tedersoo et al., 2014; Weiss et al., 2016b). Recently, it was shown that fungal derived effector molecules (called effectors) contribute to the establishment of the Sebacinales-host interactions (Nizam et al., 2019; Nostadt et al., 2020; Rafiqi et al., 2013; Wawra et al., 2016). These effectors suppress plant defense responses and modulate plant metabolism to promote compatibility in the roots, but their contribution to beneficial outcomes is unclear. Similarly, the nature of host transcriptional programs and signalling networks which lead to a mutually beneficial fungus-plant partnership are not well understood.

In the past few years, microbe-microbe interactions have emerged as an additional important element shaping plant host-microbe interactions. Using a soil-based split-root system, we demonstrated that both local and systemic colonisation by the Sebacinales endophyte *Serendipita vermifera* (syn. *Sebacina vermifera*, hereafter *Sv*) afford protection against *Bs* infection and disease symptoms in *Hordeum vulgare* (barley) (Sarkar et al., 2019). Here, we explore how *Sv* and *Bs* colonisation capacities in two plant species, barley and Arabidopsis, are modulated by the presence of individual members of the core bacterial microbiota or SynComs isolated from the barley rhizosphere (Robertson-Albertyn et al., 2021) or Arabidopsis roots (Bai et al., 2015). The finding that *Bs* also infects and causes disease symptoms in Arabidopsis roots motivated us to develop a set of physiological measurements to characterize disease severity and plant growth in Arabidopsis under different microbe treatment regimes. These measurements include ion leakage (quantified *via* electric conductivity) and photosynthetic activity (measured using pulse amplitude modulation fluorometry) as readouts for host cell death progression and biotic stress during the host-microbe interaction. Analyses of inter-kingdom activities in barley and Arabidopsis revealed that *Sv* can functionally replace root-associated bacteria by mitigating pathogen infection and disease symptoms in both hosts. Additionally, we show that cooperation between bacteria and beneficial fungi leads to inter-kingdom synergistic beneficial effects. Finally, RNA-seq experiments with selected bacterial strains alone or combined with *Sv* and/or *Bs* provide insights to how microbes synergistically protect plants. We conclude that plants have evolved to preferentially accommodate communities that support their health and that root-associated prokaryotic and eukaryotic microbes can act synergistically with the plant host in limiting fungal disease.

## Material and Methods

### Plant, fungal and bacterial materials

Barley (*Hordeum vulgare* L. cv Golden Promise) and *Arabidopsis thaliana* Col-0 were used as plant host organisms. *Serendipita vermifera* (MAFF305830) and *Bipolaris sorokiniana* (ND90Pr) were the fungal models used in this study. The *At*SynCom consists of four bacterial strains from the *At*Sphere collection (Bai et al., 2015). The *Hv*SynCom consists of 26 bacterial strains of an existing collection (Robertson-Albertyn et al., 2021) as described in Figure S1.

### Growth conditions and microbe inoculations

Barley seeds were surface sterilized in 6 % sodium hypochloride for one hour under continuous shaking and subsequently washed each 30 min for 4 h with sterile water. The seeds germinated on wet filter paper in darkness and room temperature for 4 days, transferred to 1/10 PNM (Plant Nutrition Medium, pH 5.7) (Wawra et al., 2016) in sterile glass jars for growth at a day/night cycle of 16/8 h at 22/18°C, 60 % humidity under 108 µmol m^-2^ s^-1^ light intensity post inoculation.

Arabidopsis seeds (Col-0, hereafter *At*) were surface sterilized in two times 70 % and 100 % EtOH respectively for 5 min each and sown on ½ MS (Murashige-Skoog-Medium including vitamins, pH 5.6) with 1% sucrose after ethanol removal. Following two days of stratification at 4 °C and darkness, the seeds germinated at a day/night cycle of 8/16 h at 22/18°C, 60 % humidity and a light intensity of 125 µmol m^-2^ s^-1^ for seven days. Growth matched seedlings were transferred to 1/10 PNM medium in 12x12 cm square petri dishes 1 day prior to microbe inoculation.

Single bacterial strains were grown separately in liquid TSB medium (Sigma Aldrich) (15g/ L) at 28°C in darkness shaking at 120 rpm for 1 to 3 days depending on growth rates. Final OD_600_ was adjusted to 0.01 prior to inoculation of single strains or mix in equal amounts for SynComs constitutions to a final OD_600_ of 0.01. *Sv* was propagated on MYP medium (Lahrmann et al., 2015) and *Bs* on modified CM (Sarkar et al., 2019) medium both containing 1.5% agar at 28 °C in darkness for 21 days and 14 days pre inoculation respectively. *Sv* mycelial and *Bs* conidia suspensions were prepared as described in (Sarkar et al., 2019).

Arabidopsis roots were inoculated either with *Sv* mycelium (1g/50ml), *Bs* conidia (5x10^3^ spores/ml), bacteria (OD_600_ = 0.01) or a mixture of organisms contained in 0.5ml sterile water equally spread across individual plates. Barley roots were inoculated with 3ml of *Sv* mycelium (2g/50ml), *Bs* conidia (5x10^3^ spores/ml), bacteria (OD_600_ = 0.01) or a respective mixture of organisms per jar. Sterile water was used as a control treatment. Arabidopsis and barley roots were harvested at 6 dpi. Roots of both plants were cut, washed thoroughly to remove extraradical fungal hyphae and bacteria and snap-frozen in liquid nitrogen. Per repeat of each experiment and treatments, roots from 60 Arabidopsis plants or 4 barley plants were pooled.

### Pulse-Amplitude-Modulation (PAM) fluorometry and ion leakage measurement

For PAM fluorometry and ion leakage assays, Arabidopsis seedlings were harvested at 7 dpi. The plant roots were washed carefully and thoroughly to remove extraradical fungal hyphae and bacteria and subsequently transferred to a 24 well plate containing 2 ml sterile water. Five seedlings of the same treatment were pooled in one well. PAM fluorometry and Ion leakage were measured every 24 hours for 4-7 days as previously described (Dunken et al., submitted).

### RNA isolation for RNA-seq and RT-PCR

RNA extraction for quantification of fungal colonisation and RNA-seq, cDNA generation and RT-PCR were performed as described previously (Sarkar et al., 2019). Primers used are listed in Table S1.

### Genomic and transcriptomic data analysis

Stranded mRNA-seq Libraries were prepared according to the manufacturer’s instructions (Vazyme Biotech Co., Nanjing, China). Qualified libraries were sequenced on a HiSeq 3000 system instrument at Genomics & Transcriptomics Laboratory, Heinrich-Heine University, Germany (https://www.gtl.hhu.de/en.html) to generate 50 million reads with a 150-bp read length from two to three biological replicates). Reads with Illumina adaptors and the sequence quality lower than 15 were removed using fastp (Chen et al., 2018). Reads were mapped to the annotated genomes of the three organisms (barley: IBSC Morex v2, *Bipolaris sorokiniana*: Cocsa1, *Serendipita vermifera*: *Sebacina vermifera* MAFF 305830 v1.0, Table S2). Count per gene files were generated using an in-house multi-organism mapping pipeline (Niu et al. in preparation). Read count per transcript was converted into read count per gene using R package tximport (Soneson et al., 2015). Potential batch effects were excluded with Combat-seq function in SVA package (Zhang et al., 2020). We selected 25,172 of 39,734, 10,178 of 12,250, and 13,376 of 15,312 genes having more than averaged five reads per condition for *H. vulgare*, *B. sorokiniana*, and *S. vermifera* respectively for the analysis (Table S3-5). The log2 fold difference of the gene expression between conditions was calculated with R package DESeq2 (Love et al., 2014). Genes with statistical significance were selected (FDR adjusted p value < 0.05). The consistency of normalized transcription from two to three biological replicates was confirmed by visualizing the distribution of read counts. Normalized read counts of the genes were also produced with DESeq2, which were subsequently log2 transformed. Functional annotation sets were combined using Carbohydrate Active Enzyme database (CAZy,(Lombard et al., 2014), the Gene Ontology (GO; The Gene Ontology Consortium, 2015(Gene Ontology, 2015)), Kyoto Encyclopedia of Genes and Genomes (KEGG; (Ogata et al., 1999)), and EuKaryotic Orthologous Groups (KOG;(Tatusov et al., 2003), PFAM (Finn et al., 2016), Panther (Thomas et al., 2003), MEROPS (Rawlings et al., 2018). KOG, GO, KEGG, PFAM, Panther, MEROPS, best O. sativa hit homologues, best Athaliana TAIR10 hit homologues were obtained from Phytozome, JGI (https://phytozome.jgi.doe.gov/pz/portal.html#!bulk?org=Org_Hvulgare_er). CAZymes, MEROPS, and GO terms were obtained based on KEGG, GO, PFAM, IDs using R packages KEGG.db, GO.db, and PFAM.db (Carlson, 2016; Carlson, 2019; Carlson et al., 2018). Fungal genomes and functional annotations were obtained from Mycocosm, Joint Genome Institute (https://mycocosm.jgi.doe.gov/mycocosm/home). The latest CAZy annotations were provided from CAZy team (www.cazy.org). Theoretically secreted proteins were determined with Secretome pipeline described previously (Pellegrin et al., 2015). We identified the genes coding for CAZymes, lipases, proteases, small secreted proteins (less than 300 amino acid) as a subcategory. Fungal effectors were previously identified, which were combined with the predicted secretome information in this study (Sarkar et al., 2019). We sorted significantly differentially regulated genes specific to the conditions (> 1 log2 FC; FDR adjusted p < 0.05) and visualized with R package UpSetR (Gehlenborg and Conway, 2019). Such genes were grouped using K-means clustering with R package, pheatmap (Kolde, 2019). Networks of k-means clustered genes visualised with R package, ggraph (Pedersen, 2020). Genes expressed differently among the conditions were identified based on principal coordinates calculated with R package Vegan (Oksanen et al., 2020). The first three principal coordinates were used to select high loading genes coding for glycosyl hydrolases and effectors of *B. sorokiniana*. Comparative analyses with a previous transcriptomic dataset (Sarkar et al., 2019) showed that 37 of the 50 top induced barley genes in response to *Bs* in soil are again detected to be significantly induced in the Barley_*Bs* vs Barley comparison in PNM (this study), indicating a large overlap of the highly responsive host genes to the pathogen in soil and PNM. Data are deposited to the NCBI under the BioProject accession number: PRJNA715112.

### Gene co-expression analysis

A self-organizing map (SOM) was trained with the normalized read count of the selected replicates using Rsomoclu and kohonen (Peter Wittek, 2017; Ron Wehrens, 2007). The total of 1015 nodes (35 x 29 matrix was used with a rectangular shape (four neighbouring nodes). The resolution of 25 genes per node was applied for clustering, which was empirically optimised (Miyauchi et al., 2016; Miyauchi et al., 2017). The epoch of 1000 times more than the map size was applied (i.e., 1,015,000 iterations of learning, being 1015 map size times 1000). The genes showing similar regulation trends were grouped based on the mean transcription of the nodes. We examined genome-wide condition-specific transcriptomic patterns in graphical outputs (i.e. Tatami maps). Mean transcription values were calculated from the grouped genes per condition in each node (i.e. node-wise transcription). Then, using the node-wise transcription values, highly-regulated genes specific to each of the conditions were determined by fulfilling either of two criteria: 1) > 12.6 log2 reads (above 95th percentile of the entire transcribed genes); or 2) over ± 2 log2 transcriptional differences between testing conditions and a control. The process above was performed in a semi-automated manner using co-gene expression pipeline (SHIN+GO; (Miyauchi et al., 2020; Miyauchi et al., 2016; Miyauchi et al., 2017; Miyauchi et al., 2018). R was used for operating the pipeline (R Core Team, 2013).

## Results

### Sebacinales associate with healthy Arabidopsis plants in diverse European locations

By monitoring root-associated microbial communities in natural *A. thaliana* populations, Thiergart et al. (Thiergart et al., 2020b) showed that microbial community differentiation in the roots is explained primarily by location for filamentous eukaryotes and by soil origin for bacteria, whereas host genotype effects are marginal. We re-analysed this dataset, including lower abundance operational taxonomic units (OTUs), and found that fungal OTUs of the order Sebacinales were significantly enriched in the rhizoplane compartment of healthy Arabidopsis plants in diverse European locations (Figure 1). These environmental sampling data complement cytological studies which show that Sebacinales isolates colonize Arabidopsis by forming a loose hyphal mesh around roots with intracellular colonisation limited to the root epidermis and cortex layer (Lahrmann et al., 2015). The frequent occurrence and enrichment patterns of Sebacinales OTUs in the roots of native Arabidopsis suggest a functional endophytic association with this host in nature. This finding motivated us to investigate the functional relevance and resilience of these fungi in a community context in the roots of Arabidopsis and to compare these with the beneficial effects observed in barley using bacterial synthetic communities.

**Fig. 1:**
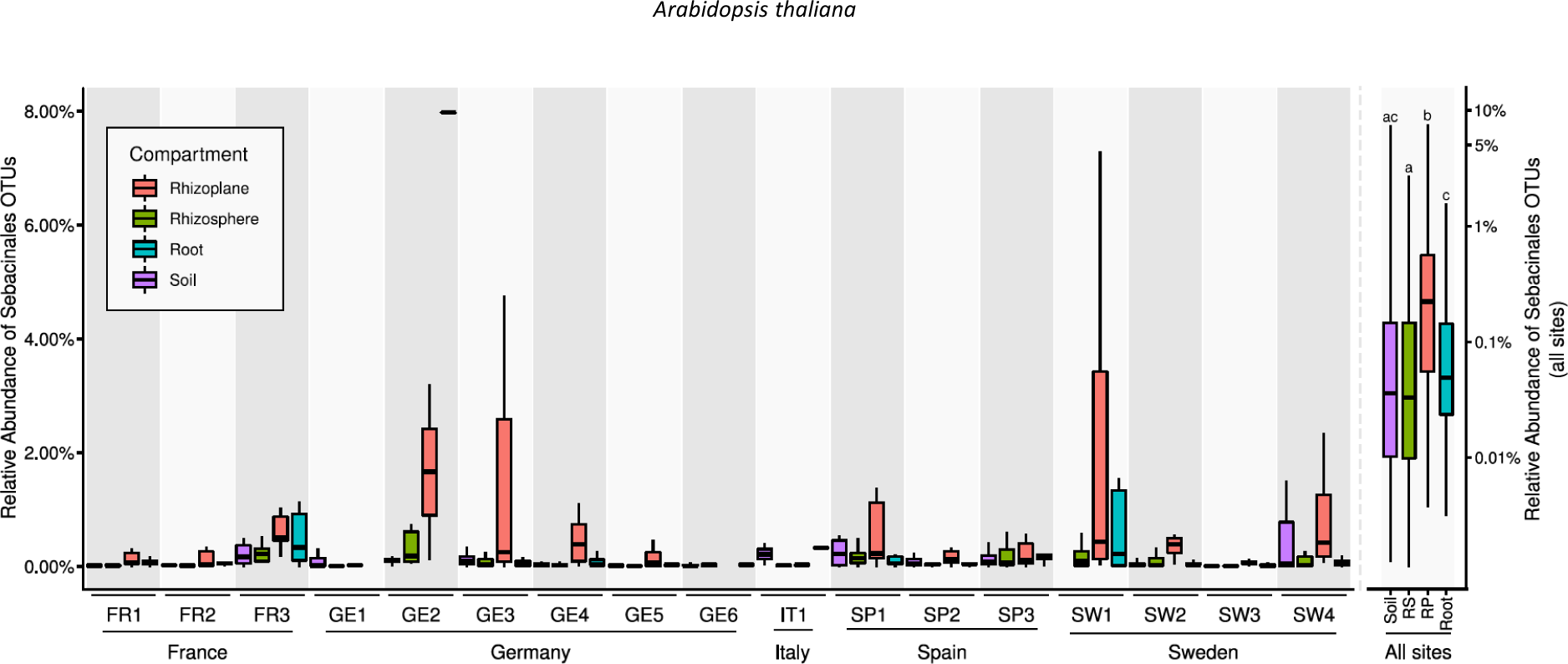
Abundance of Sebacinales in Arabidopsis roots of different European locations. A) Analysis of fungal (ITS1) OTUs belonging to the Sebacinales order from sequencing data obtained from samples of soil and root-associated microbial communities across 3 years and 17 European sites where naturally occurring *A. thaliana* populations were found (Thiergart et al., 2020). A non-parametric Kruskal-Wallis test with a Dunn’s test for multiple comparisons on relative abundances of Sebacinales OTUs in different compartments, aggregated for all site, shows that this fungal taxon is enriched in the rhizoplane compartment of *A. thaliana* roots compared to the other compartments.

### Protection mediated by *S. vermifera* and bacteria is synergistic and largely independent of the host

We reported that *Sv* acts as an extended plant protection barrier in the rhizosphere which reduces barley root infection and disease symptoms caused by the hemibiotrophic pathogen *Bs* on defined plant sugar-free minimal medium (PNM) and in natural soil (Sarkar et al., 2019). Here we confirmed the protective activity of *Sv* during *Bs* infection of barley root tissue (Figures 2A-D) and additionally we observed enhanced *Sv* colonization through the presence of *Bs* at 6 days post inoculation (dpi) on PNM (Figure 2B).

**Fig. 2:**
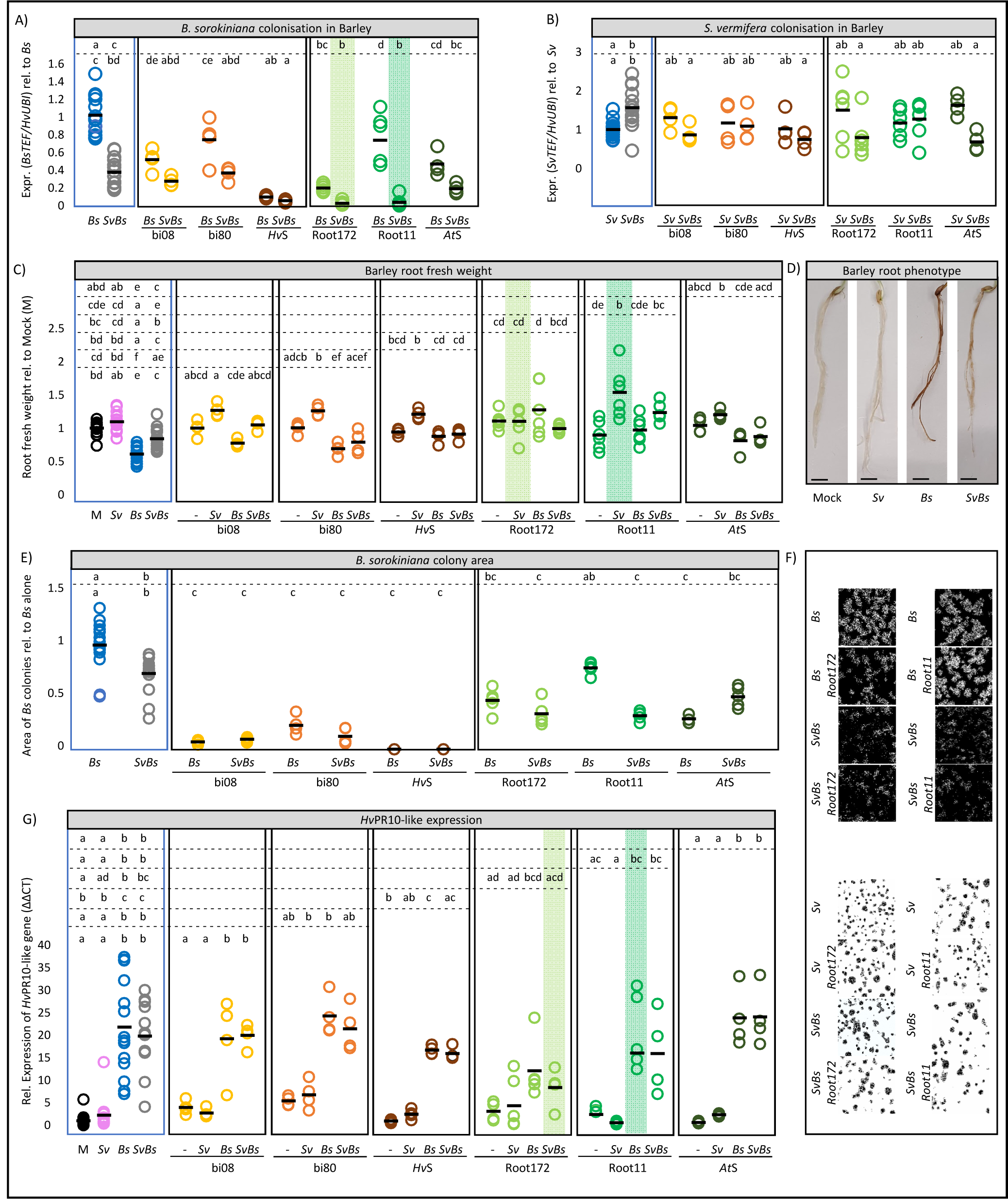
Barley root colonisation and responses after fungal and/or bacterial inoculation at 6 dpi. A) *Bs* and B) *Sv* colonisation in barley at 6 d post inoculation. Fungal colonisation in each biological replicate was confirmed by quantitative RT-PCR inferred by expression analysis of the fungal housekeeping gene TEF compared with barley ubiquitin (UBI) gene (n = 4-14). C) Barley root fresh weight per biological replicate normalised to water (Mock) inoculated plants (n = 4-14 with 4 plants each). D) Pictures showing barley roots inoculated with water as a control (Mock), *Sv*, *Bs* or both fungi, scale bar = 1 cm). E) *Bs* colony area in direct confrontation with *Sv* or bacteria in absence of the host on defined medium relative to *Bs* alone. F) Pictures of direct confrontation assays. *Bs* colonies (black background) and *Sv* colonies (white background) were filtered using ImageJ and the morphoLipJ plugin. *Sv* colony area was not negatively affected by the presence of the other microbes (data not shown). G) Relative expression of HvPr10-like gene (HORVU0Hr1G011720). Green background highlights samples that were later used for RNAseq. Different letters in the comparison between the tripartite panel (blue square) and combinations of any other panel (defined by the dashed lines) represent statistically significant differences according to one-way ANOVA and Tukey‘s post-hoc test (p < 0.05).

To establish whether *Sv* antagonizes root infection by *Bs* in other plant hosts, we assessed fungal colonisation and disease symptoms in root tissues of Arabidopsis with *Sv*, *Bs* or both fungi on PNM (Figure 3A). *Bs* infected Arabidopsis seedlings displayed prominent disease symptoms at 6 dpi such as reduced main root length, rosette diameter and lateral root number compared to mock inoculated controls (Figures 3B, S2B and S2C). *Bs* inoculated roots exhibited characteristic tissue browning (Figure S2G), increased ion leakage and a reduced photosynthetic active leaf area over time, indicative of host cell death progression (Figures 3E-I and S2F). As shown for barley and in accordance with their growth rates in axenic cultures (Sarkar et al., 2019), *Bs* generated more endophytic biomass than *Sv* upon separate inoculations of Arabidopsis roots, determined by a quantitative reverse transcription PCR (RT-qPCR) test displaying the ratio between constitutively expressed single copy fungal (TEF) and plant (UBI) genes (Figures 3C and 3D). Notably, *Bs* endophytic biomass and disease symptoms were substantially diminished in roots that were co-colonized by *Sv* (Figures 3B, 3C and S2). In contrast, *Sv* endophytic colonisation was enhanced by the presence of the pathogen also in this tripartite interaction (Figure 3D). The enhanced *Sv* colonization in both hosts could be explained by the plant actively recruiting *Sv* to suppress the soil-born pathogen or *Sv* feeding on *Bs* and/or necrotic plant tissues.

**Fig. 3:**
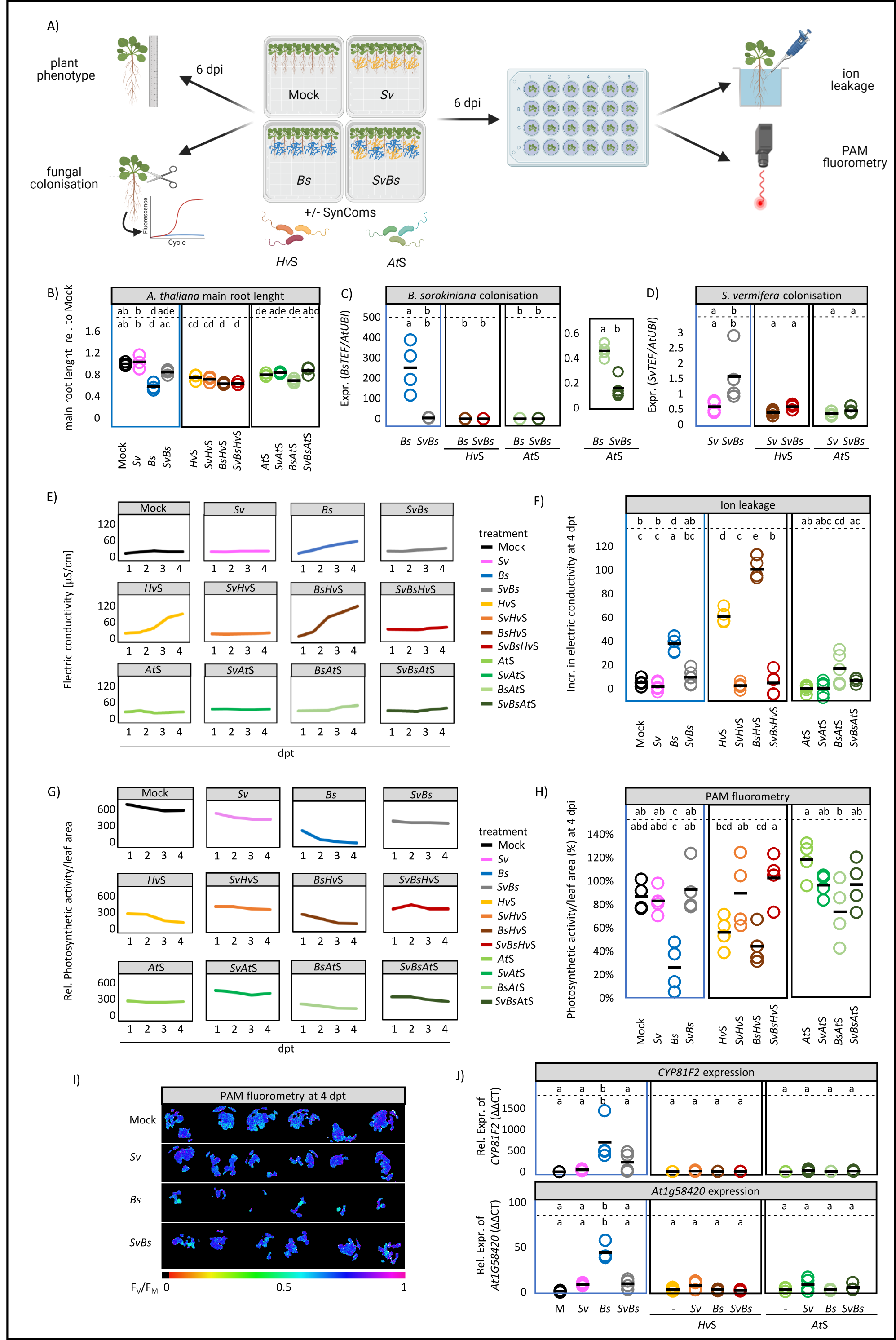
Arabidopsis root colonisation and responses after fungal and/or bacterial inoculation at 6 dpi. A) schematic drawing of the experimental setup measuring the electric conductivity (ion leakage) and photosynthetic activity (PAM fluorometry) in Arabidopsis seedlings. B) the main root length of *A. thaliana* inoculated in dipartite, tripartite and multipartite systems with *B. sorokiniana* (*Bs*), *S. vermifera* (*Sv*) and the bacterial synthetic communities *Hv* SynCom (*Hv*S) or *At* SynCom (*At*S). C) *Bs* and D) *Sv* colonisation in *A. thaliana* at 6 d post inoculation inferred by expression analysis of the fungal housekeeping gene TEF compared with Arabidopsis ubiquitin (UBI) (n = 4). To further assess *Bs* disease symptoms and plant health we measured E) the electric conductivity from 1 – 4 d post transfer (n = 6); F) the total increase in electric conductivity from 1 – 4 d post transfer (n = 6); G) the photosynthetic activity (F_V_/F_M_) from 1 – 4 d post transfer (n = 6); and H) the photosynthetic activity per leaf area at 4 dpi relative to 1 dpi (n = 6). I) The photosystem II (PSII) quantum yield of 5 At seedlings/well at 4 dpt after dark adaptation (F_V_/F_M_) via PAM flourometry. Purple/dark blue, lighter colors and black color indicate high, reduced and lack of PS II activity respectively. *Bs* infection continuously reduces local PS II activity and spreads across the whole seedling, leading to a reduced photosynthetic active leaf area over time. J) The expression of the fungal responsive gene At1g58420 and the gene encoding for the cytochrome P450 monooxygenase CYP81F2 involved in indole glucosinolate biosynthesis and defense. Statistical analyses were performed for each subpanel together with the tripartite panel (in blue). Different letters in the comparison between the tripartite panel (blue square) and combinations of any other panel (defined by the dashed lines) represent statistically significant differences according to one-way ANOVA and Tukey‘s post-hoc test (p < 0.05).

Next, we determined whether bacterial strains isolated from the rhizosphere of barley (*Hv*SynCom) or the endosphere of Arabidopsis roots (*At*SynCom) can also protect barley and Arabidopsis from *Bs* infection. Both SynComs were able to reduce *Bs* colonisation and largely rescue plant growth phenotypes caused by the pathogen in both hosts (Figures 2A, 2C, 3B, 3C and S2). Interestingly, the *Hv*SynCom alone, but not the *At*SynCom, caused increased ion leakage and reduced photosynthetic active leaf area in Arabidopsis (Figures 3E-H and S2A). This points towards an induction of host cell death in Arabidopsis by the non-native bacterial SynCom.

To clarify whether the observed host protection against *Bs* infection is a general property of root-associated bacterial strains or requires a community context, we inoculated functionally and taxonomically-paired bacterial strains from the *Hv-* and *At-*SynComs (Figure S1) individually or in combination with *Bs* on barley. We observed a strong reduction of the pathogen infection with the Proteobacteria strains bi08 (*Pseudomonas* sp.) and Root172 (*Mesorhizobium* sp.) but not with the Firmicutes strain bi80 (*Bacillus* sp.) and only marginally with Root11 (*Bacillus* sp.) irrespective of the host species origin (Figure 2A). This indicates that not all bacterial strains in the SynComs have the ability to protect the roots from *Bs* infection but the overall protection effect is maintained in a community context.

Next, we interrogated whether the observed beneficial effects on the plant hosts mediated by *Sv* or the bacterial strains are retained or altered during inter-kingdom interactions. For this, we co-inoculated barley and Arabidopsis roots with *Sv* and *Bs* in combination with a single bacterial strain or the SynComs. We found that *Sv* colonisation was only marginally affected by the presence of the bacteria (Figures 2B and 3D). The combined presence of *Sv* and bacterial strains led to a stabilized (reduced biological variation) or potentiated host protection against *Bs* infection (Figure 2A, 3C and S2). Potentiated protection to *Bs* infection was most evident during co-inoculation of *Sv* with Root11 in barley (Figures 2A and 2C). These data show a robust inter-kingdom protective effect of *Sv* with bacteria against an invasive fungal root pathogen.

Finally, to measure whether the host plant contributes to the effects displayed by *Sv* and the examined bacterial strains in limiting pathogen biomass, we additionally performed direct microbe-microbe confrontation assays on PNM. In these assays we largely recapitulated the antagonism observed against *Bs in planta* (Figure 2E and 2F). We therefore concluded that microbe-microbe interactions rather than the host plant are most important for conferring the root protective properties of *Sv* or the tested bacteria. This notion is also supported by *in planta* cytological analyses in which we observed a direct interaction between *Bs* and Root172 at the rhizoplane of Arabidopsis and extensive lysis of the fungal extracellular polysaccharide matrix surrounding *Bs* hyphae (Figure 4).

**Fig. 4:**
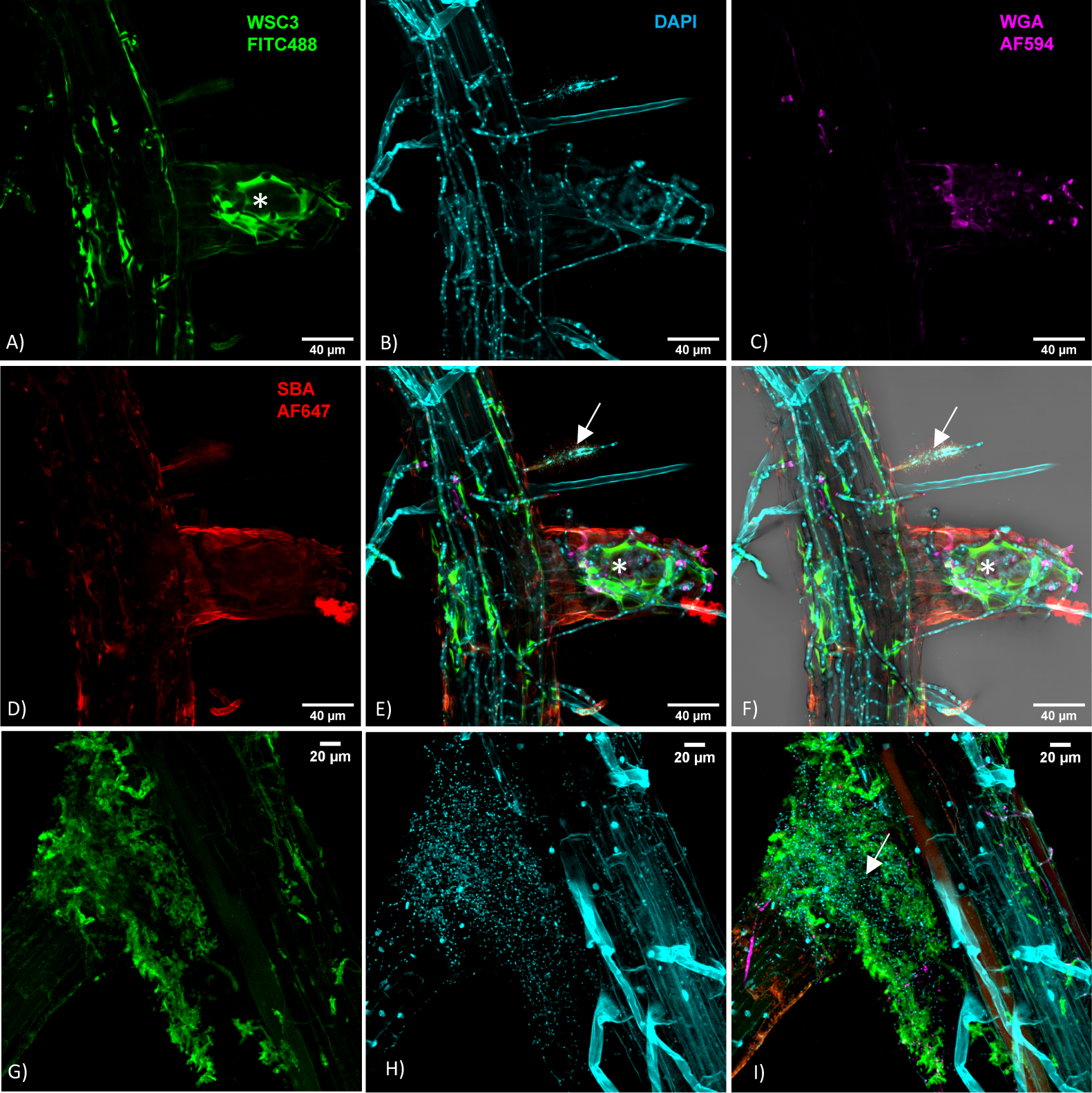
*Arabidopsis thaliana* Col-0 inoculated with *Bs*+Root172 at 7 dpi. Roots were fixed with 70% EtOH and stained with the β-1,3-glucan binding lectin WSC3-FITC488 which binds to the fungal matrix (in A and G), the fluorescent DNA stain DAPI (in B and H), the chitin stain WGA-AF594 (in C) and the lectin SBA AF647, which binds α- and β-N-acetylgalactosamine and galactopyranosyl residues (in D). Overlay in E, F and I. White arrows: *Bs* hyphae after loss of matrix in the presence of Root172. Asterisks: intact fungal matrix.

### *S. vermifera* confers plant growth promotion in cooperation with selected root-associated bacteria

*Sv* promotes plant growth in different host species at late stages of colonisation (Barazani et al., 2005; Ghimire et al., 2009; Waller et al., 2008). At an early colonisation time point of 6 dpi in barley, neither *Sv* alone nor any of the single bacterial strains or SynComs led to a significant change in root fresh weight (Figure 2C). By contrast, a combination of *Sv* and bacterial strains Root11, bi08 or bi80, significantly increased barley root fresh weight at 6 dpi (Figure 2C). This early inter-kingdom mediated root growth promotion effect was strain-specific, not restricted to bacterial strains isolated from the barley rhizosphere, and maintained in a community context. Co-inoculation with heat-inactivated bacterial SynComs failed to increase barley root fresh weight (Figure S3), underlying the importance of living bacteria in promoting root growth.

In Arabidopsis, we observed root growth inhibition at 6 dpi upon inoculation with *Bs* or the SynComs irrespective of the number of bacterial strains and their host origin (Figure 3B). Co-inoculation with *Sv* largely alleviated the *Bs*-mediated root growth inhibition but did not increase root or shoot size compared to controls (Figures 3B, S2B and S2C). Only the combination of Root172 with *Sv* led to a significant increase in Arabidopsis rosette diameter at 6 dpi (Figures S2D and S2E). This phenotype was, however, not retained in a bacterial community context, suggesting that it is less robust and/or plant growth promoting microbes suffer from competition by other community members. Put together, our data suggest that the establishment of beneficial inter-kingdom interactions in the plant microbiota is an evolutionarily conserved trait that can be fine-tuned by bacterial composition and host species.

### Inter-kingdom synergistic beneficial activities are not associated with extensive host transcriptional responses

To investigate mechanisms underlying the synergistic beneficial effects displayed by a combined fungal endophyte and bacterial inoculation, we analysed the barley root transcriptome during fungal and bacterial colonisation by RNA-seq. The multipartite systems used for transcriptomics included the two fungi (*Sv* and *Bs*) and the bacterial strains Root172 or Root11, selected based on their distinctive and robust *in planta* activities with *Bs* and *Sv* at 6 dpi. Namely, Root172 conferred strong host protection against *Bs* whereas Root11 had a strong root growth promotion phenotype (Figures 2A and 2B). To determine species representation in the Illumina RNA-seq reads, we mapped reads to annotated genes of the barley and fungal reference genomes. Bacterial reads were not present in the dataset due to the method used for the library preparation. On average, 7.9% of reads matched *Sv* genes in all endophyte-containing samples (Figure 5A; Table S2). By contrast, the relative abundance of reads mapping to *Bs* genes decreased from 13.1% (*Bs* alone) to 8.6%, 12% or 5% when *Sv*, Root11 or Root172 were co-inoculated with the fungal pathogen, respectively. Co-inoculation of Root11 or Root172 with *Sv* and *Bs* reduced the relative abundance of pathogen reads, to 2.6% and 2.7%, respectively. The reduction in *Bs* reads with *Sv* and/or bacterial strains likely reflects reduced *Bs* biomass, confirming the quantitative RT-PCR analysis (Figure 2A). To dissect barley transcriptomic trends and identify differentially expressed genes (DEG), we examined genes that were induced or repressed under specific conditions after transcript mapping and quality assessment (Figure S5, see Methods). Consistent with our previous data (Sarkar et al., 2019), we detected only a weak host transcriptomic response to *Sv* (184 DEG with log2FC>1, Figure 5C; Table S7). Neither presence of the bacterial strains nor combined presence of bacteria and *Sv* led to an extensive host transcriptional response (Figures 5C and S6; Table S7). Thus, the observed early root growth promoting effects mediated by *Sv* with Root11 in barley were not accompanied by a strong host transcriptional response (with 13 DEG specific to this condition, Figures 5C and S6; Table S7).

**Fig. 5:**
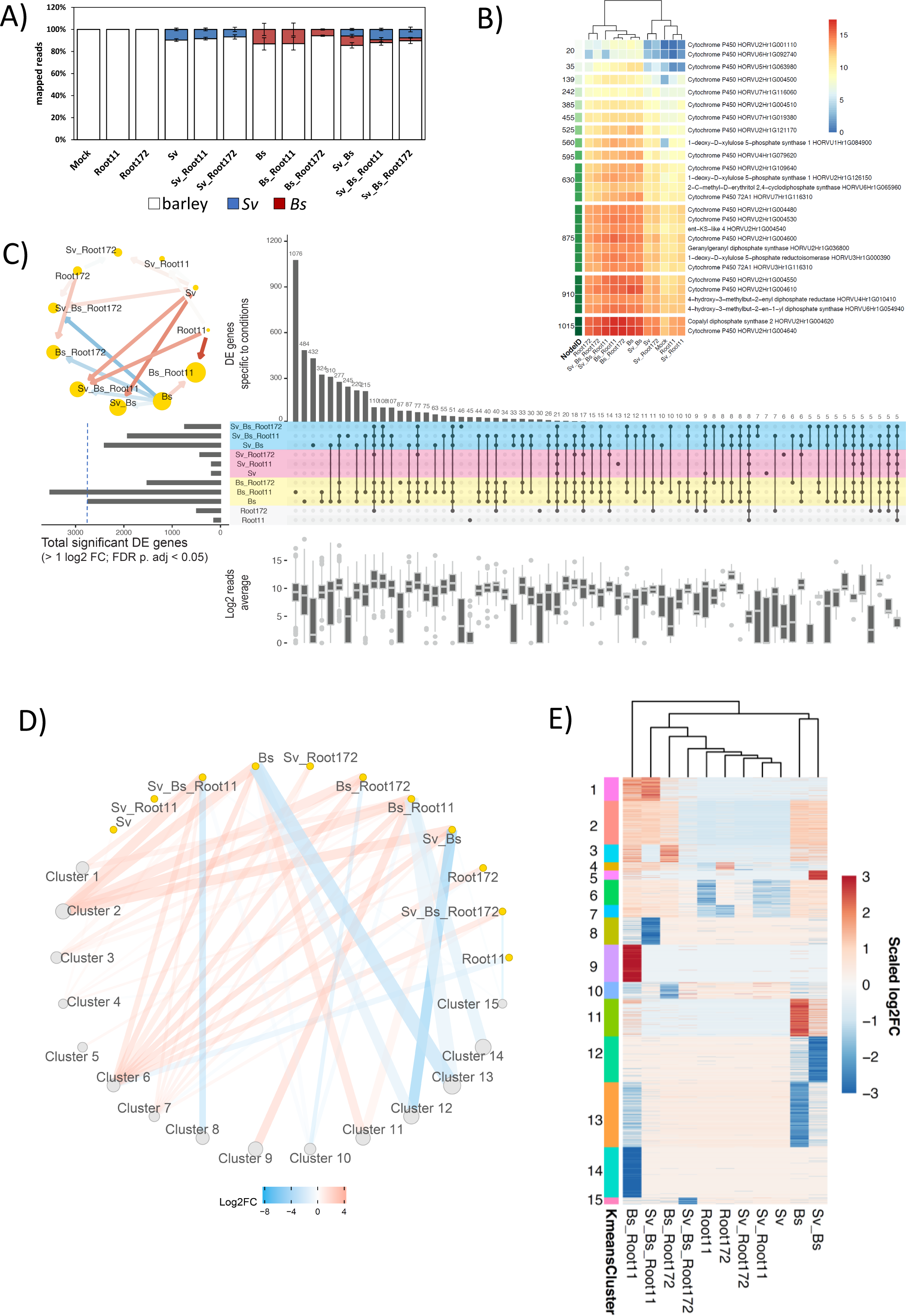
Analysis of barley root transcriptional responses to fungal and bacterial inoculation at 6 dpi. *Bs*: *Bipolaris sorokiniana*. *Sv*: *Serendipita vermifera*. Root11 & Root172: *A. thaliana* root-associated bacterial strains Root11 & Root172. A) Proportion of reads mapped to the organisms per sample. A total of 34 RNA-seq samples were mapped to the corresponding organisms. Mock: *Hordeum vulgare*. See Table S2. B) Transcription level of genes putatively involved in terpenoid phytoalexin synthesis. Averaged transcription in log2 is shown per condition. Terpenoid phytoalexin synthesis pathway in barley was published earlier (Sarkar et al., 2019). See Tab. S6. C) Condition-specific differentially expressed genes (> 1 log2FC; FDR adjusted p-value < 0.05) are compared to barley Mock control. Horizontal bars: Total number of DE genes per condition, additionally visualised as a network. Red and blue arrows represent a normalised high and low number of DEGs among the conditions compared. The size of the circles corresponds to the total number of DEGs. Vertical bars: Number of genes unique/shared for top 70 intersections. See Table S7. D) K-means clustering of differentially expressed genes grouped into 15 clusters visualised as a network. Node size and line thickness correspond to the number of DEGs. Colours of lines connecting clusters and conditions represent log2 fold changes and up/down regulations. E) K-means clustering of above is presented as a heatmap. A total of 14,274 differentially expressed genes are used for C, D, and E. See Tab. S7 and Tab. S8.

**Fig. 6:**
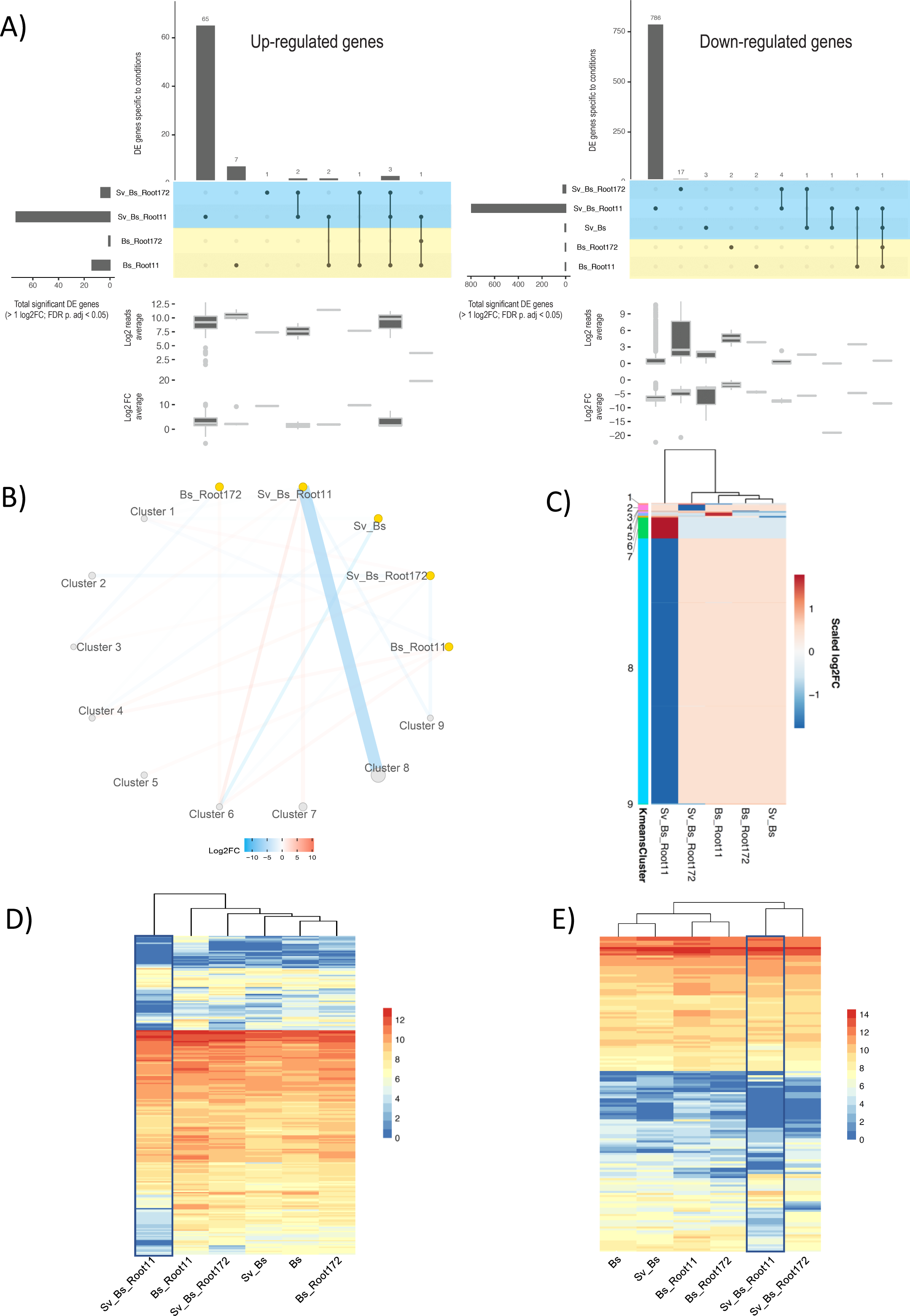
*B. sorokinana* transcriptional responses to *S. vermifera* and bacteria at 6 dpi during colonisation of barley host plants. *Bs*: *Bipolaris sorokiniana*. *Sv*: *Serendipita vermifera*. Root11 & Root172: *A. thaliana* root-associated bacterial strains Root11 & Root172. A) Condition-specific differentially expressed *B. sorokiniana* genes (> 1 log2FC; FDR adjusted p-value < 0.05) compared to barley infection alone. Horizontal bars: Total number of DEGs per condition. Vertical bars: Number of genes unique/shared for intersections. See Tab S7. B) K-means clustering of differentially expressed genes grouped into 9 clusters visualised as a network. Node size and line thickness correspond to the number of DEGs. Colours of lines connecting clusters and conditions represent log2 fold changes and up/down regulations. C) K-means clustering of the 9 groups is displayed as a heatmap. A total of 923 differentially expressed genes are used for B and C. See Tab. S8. D) Averaged log2 read counts of predicted secreted CAZyme coding genes. E) Averaged log2 read count of effector coding genes. See Tab. S9.

Conversely, infection with *Bs* resulted in 2,743 barley DEG. Co-inoculation of *Bs* and Root172 reduced barley DEG to 1,517, whereas Root11 with *Bs* produced a larger number of DEG (3,528) compared to *Bs* alone (Figures 5C and S6). Grouping DEG according to expression patterns identified 15 clusters of highly up or down regulated barley genes specific to one or more condition/s (Figures 5D and 5E; Table S8) and showed that the barley response to co-inoculation with *Bs* and Root11 was most different from all other conditions (Figure 5E). To identify functional categories in co-regulated genes, we employed a self-organizing map (SOM) to group genes into nodes displaying similar regulation (Figure S7; Table S6) and we performed GO enrichment analyses (Figure S8). These analyses showed that *Bs* alone strongly induced a barley immune response and terpenoid phytoalexin production. Root11 had no effect on immunity or terpenoid phytoalexin production, whereas Root172 slightly induced an immune response. Notably, co-inoculation of Root11 with *Bs* provoked a higher activation of immunity genes and repression of host cell wall biosynthesis and DNA modification compared to the pathogen alone (Figure S8; Table S8).

In accordance with the reduction of *Bs* biomass and disease symptoms, the presence of *Sv* reduced the number of barley DEGs in response to *Bs* (Sv_Bs: 2,403). This reduction was most pronounced in combination with the bacterial strains, especially with Root172 which had the strongest effect on *Bs* colonisation (Sv_Bs_Root11: 1,921; Sv_Bs_Root172: 740; Figures 5C and S6; Table S8). Consistently, the expression of barley genes associated with terpenoid phytoalexin production was partially reduced in the multipartite interactions compared to *Bs* alone (Figure 5B). The barley root gene expression data shows that the cooperative action of *Sv* with bacteria protect barley roots from *Bs* infection without extensive host transcriptional mobilization of immunity and defense metabolic pathways.

To test the above observation further, we investigated the immune modulatory proprieties of the beneficial *Sv* fungal and bacterial strains in roots of Arabidopsis and barley by using specific marker genes. In Arabidopsis, we observed a reduction of the expression of the gene encoding for the cytochrome P450 monooxygenase CYP81F2 involved in indole glucosinolate biosynthesis and defense (Pfalz et al., 2009) in *Bs* infected roots co-inoculated with *Sv* and/or the bacteria compared to *Bs* alone (Figure 3J). Similarly, the MAMP (microbe-associated molecular pattern) and fungal-responsive *At*1g58420 gene (Nizam et al., 2019) displayed lower expression during the multipartite interactions (Figure 3J), suggesting a reduced host response to *Bs* which correlates well with the pathogen load. In barley, we previously identified a *PR10* family gene (HORVU0Hr1G011720, hereafter referred to as *HvPR10*-like) as a robust marker for induced immune responses to *Bs* colonisation (Sarkar et al., 2019). RNA-seq and quantitative RT-PCR analyses confirmed that *HvPR10*-like expression was highly induced by *Bs* infection of barley roots. By contrast, *HvPR10*-like expression was weakly induced by *Sv* and/or the bacterial strains (Figure 2G). Despite the strong reduction in pathogen infection and disease symptoms upon co-inoculation with *Sv* and bacteria, we found that *Bs*-induced *HvPR10-like* expression was generally maintained in all combinations (Figure 2G). This result indicates that *HvPR10-like* expression is driven principally by the pathogen and impacted less by the presence of *Sv* and bacteria. Only co-inoculation of Root172 and *Sv*, which displayed the strongest protection against *Bs* infection, significantly lowered *Bs*-induced *HvPR10-like* gene expression. Hence, in conclusion, despite the general decreased barley transcriptional response to *Bs* and the lower pathogen load, the activation of specific immune responses such as the *HvPR10-like* gene were still in place in the presence of *Sv* and/or bacteria.

### Synergistic actions of *S. vermifera* and bacteria reduce the virulence potential of endophytic *B. sorokiniana*

To examine mechanisms underlying the cooperative antagonistic behaviour of *Sv* and the bacteria towards *Bs*, we analysed the fungal transcriptomes during barley root colonisation at 6 dpi. We previously reported that fungal transcriptome changes are driven mainly by their interactions with the host and that *Sv* effects on the *Bs* transcriptome occur mostly in the rhizosphere (Sarkar et al., 2019). Consistent with this notion, *Sv* or the bacterial treatments alone had little impact on the transcriptome of endophytic *Bs.* By contrast, the combined presence of *Sv* and Root11 had a strong impact on the *Bs* transcriptome with 65 up- and 786 down-regulated genes (Figure 6A; Table S8). DEG of *Bs* during root infection were grouped into nine clusters (Figures 6B and 6C; Table S8). The largest *Bs* cluster (#8) contained genes that were repressed compared to *Bs* infection of barley alone. Among the top 10 repressed genes in this cluster there were 4 *Bs* genes encoding for glycoside hydrolases (Table S8). This prompted us to look into the expression of all *Bs* CAZyme and effector genes.

We observed a general repression for these categories by the combined presence of *Sv* and Root11, possibly explaining the reduced *Bs* colonisation of roots (Figures 6D, 6E, S9 and S10; Table S9). Notably, *Bs* gene cluster #7 (with genes specifically induced in the combined presence of *Sv*_*Bs*_Root11, Figure S13; Table S10) (Heine et al., 2018; Ola et al., 2014; Zhen et al., 2018) contained six up-regulated genes potentially participating in the production of antibacterial compounds related to chrysoxanthone, neosartorin and emodin. Hence, it is possible that *Bs* actively engages in antagonizing Root11 in the presence of *Sv* at 6 dpi. On the other hand, upon *Bs* co-inoculation with Root11 we observed induced expression of fungal effector and CAZyme genes (Figures 6D, 6E, S9, S10 and cluster 5 in Figure 6C) such as several AA9, GH43, CE5, PL1 and PL3 that are known to be enriched in plant associated fungi (Lahrmann et al., 2015; Zuccaro et al., 2011), which might explain the increased host immune response in this interaction. Transcriptional changes in endophytic *Sv* in response to the other microbes in barley roots were generally smaller and predominantly driven by *Bs* pathogen load and the associated barley immune response (Figure 7, S11 and S12; Table S7-9). This is in agreement with our previous data which suggests that *Sv* transcriptional response is likely driven by the changes in the plant host environment due to the pathogen activity rather than by direct interaction with *Bs* inside the root (Sarkar et al., 2019).

**Fig. 7:**
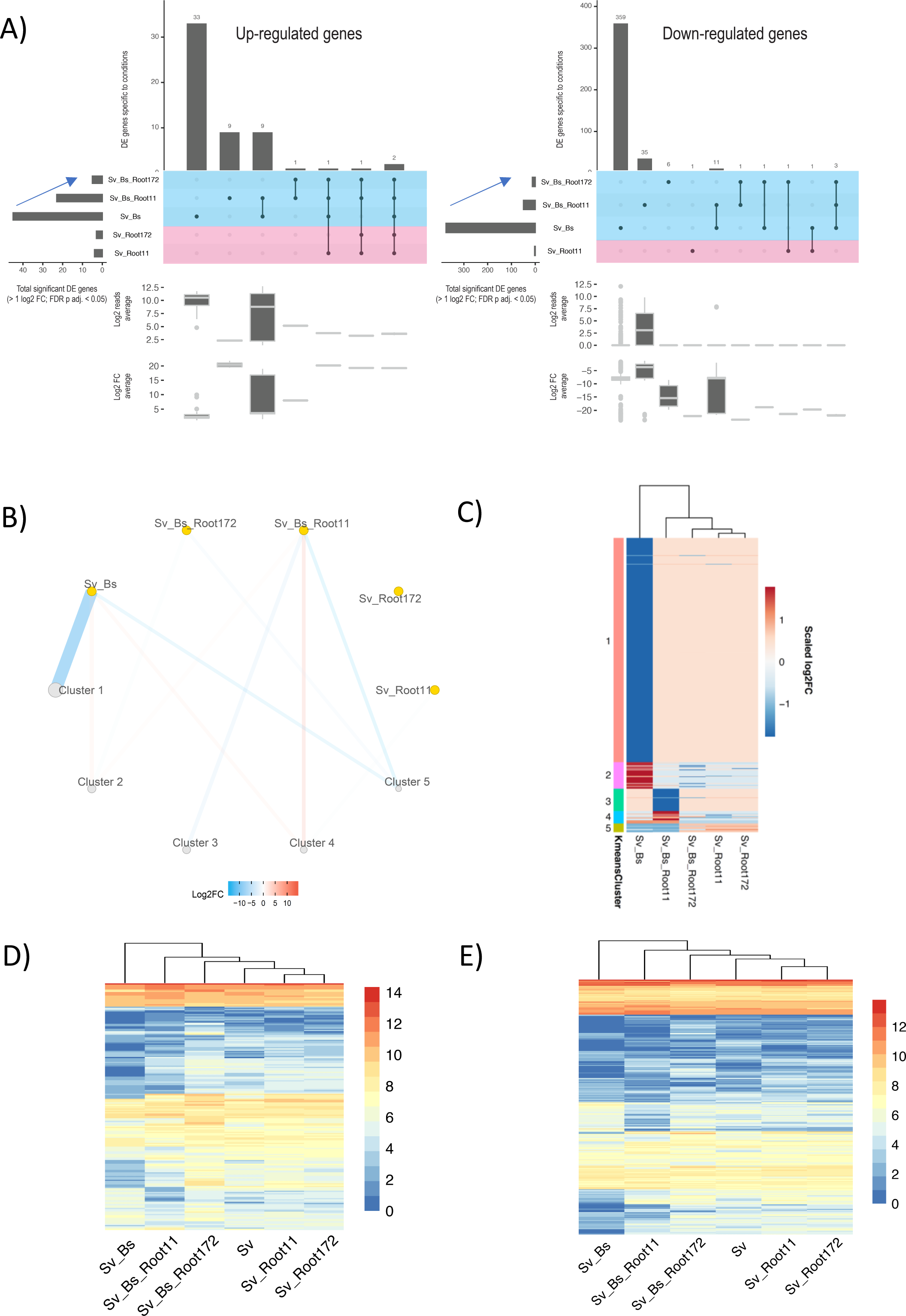
*S. vermifera* transcriptional responses to *B. sorokinana* and bacteria at 6 dpi during colonisation of barley host plants. *Bs*: *Bipolaris sorokiniana*. *Sv*: *Serendipita vermifera*. Root11 & Root172: *A. thaliana* root-associated bacterial strains Root11 & Root172. A) Condition-specific of differentially expressed genes (> 1 log2FC; FDR adjusted p-value < 0.05) are identified by comparing to the control condition (i.e. fungus alone). Horizontal bars: Total number of DEGs per condition. Vertical bars: Number of genes unique/shared for intersections. See Tab. S7. B) K-means clustering of differentially expressed genes forming 5 clusters is displayed as a network. Node size and line thickness correspond to the number of DE genes. Colours of lines connecting clusters and conditions represent log2 fold changes and up/down regulations. C) K-means clustering above is given as a heatmap. A total of 520 differentially expressed genes are used for B and C. See Tab. S8. D) Averaged log2 read count of predicted secreted CAZyme coding genes. E) Averaged log2 read count of effector coding genes. See Tab. S9.

## Discussion

In complex environments, plant-microbe interactions are not only shaped by the plant immune system (Dangl and Jones, 2001; Jones and Dangl, 2006; Pieterse et al., 2014) but also by microbe-microbe competition and co-operation, acting directly on or as an extension to plant immunity (Card et al., 2016; Snelders et al., 2018). Recent studies reveal the importance of root associated bacteria for plant survival and protection against fungi and oomycetes (Bulgarelli et al., 2013; Cha et al., 2016; Duran et al., 2018; Mendes et al., 2011; Santhanam et al., 2015). Much less attention has been paid to the role of widely distributed beneficial endophytic fungi in a multi-kingdom context. Here we show that the effects on host growth and protection that are conferred by the Sebacinales member *S. vermifera* in bipartite and tripartite interactions (Deshmukh et al., 2006; Sarkar et al., 2019) are retained in a community context. The observed robust protective function and stability of *Sv* colonisation is likely due to its ability to adapt to changes in the plant host environment (Sarkar et al., 2019). The strength of its protection against an aggressive root fungal pathogen (*Bs*) is underscored by the observation that *Sv* can functionally replace core bacterial microbiota members in mitigating pathogen infection and disease symptoms in distantly related plant hosts. This finding is in accordance with Arabidopsis root microbiota samplings across European habitats which shows Sebacinales fungi to be of low abundance but consistently present in the host roots and the rhizosphere. Our data highlight the potential importance of less abundant but widespread root fungal endophytes in maintaining plant host physiological fitness in nature, thereby emphasizing that numerically inconspicuous microbes can play a significant role in microbiota functional studies and should be considered when designing SynComs with multiple traits, such as resilience and specific beneficial functions.

Strikingly, the presence of *Sv* also reduced the negative effect caused by the *Hv*SynCom in Arabidopsis (Figures 3E-J), revealing a more general protective activity of root endophytic fungi. The induction of cell death by the barley derived SynCom in Arabidopsis could be due to the presence of specific bacterial strains that are absent in the *At*SynCom. One such bacterial group that is well represented in the *Hv*SynCom but absent in the *At*SynCom used in this study is the Pseudomonadales. Several members of this group are reported to be pathogenic (Xin et al., 2018) whereas others with very few genome differences promote plant growth and exert biocontrol activities against different fungal pathogens (Mercado-Blanco and Bakker, 2007). However, we did not observe an increase in ion leakage upon inoculation with the *Pseudomonas* strain bi08 or other members of the *Hv*SynCom when inoculated alone (Figure S4). The pathogenicity of a single bacterial strain is likely to be suppressed in a community context, as observed for *Bs* (Figures 2 and 3). Thus, another explanation to the negative effects of the *Hv*SynCom in Arabidopsis but not in barley might be a lack of adaptation to Arabidopsis. This notion is supported by a recent analysis which detected a clear signature of host preferences among commensal bacteria from diverse taxonomic groups, including Pseudomonadales in Arabidopsis and *Lotus japonicus* (Wippel et al., 2021).

Our transcriptomic analyses show that effects of the tested bacterial strains in tripartite associations differ substantially. The general decreased barley transcriptional response to the pathogen driven by the Rhizobiales strain Root172 (Figure 5C) and the lysis of the fungal matrix at the host rhizoplane suggest that this bacterial strain act mostly directly on *Bs* (Figure 4). This is also supported by the strong antagonism of *Bs* growth irrespective of the presence of a host plant (Figures 2A, 2E and 2F). Taken together, these results point to Root172 as a possible biocontrol agent against *Bs* and potentially other root-infecting pathogens. The impact of Root172 contrasted strikingly with that of the Bacillales strain Root11 which did not limit *Bs* growth but rather enhanced *Bs* pathogenicity in barley. Notably, combining these two bacterial strains with *Sv* led to a restriction of *Bs* that exceeded the protective benefits of *Sv* and the bacteria alone (Figure 3C). These synergistic beneficial effects are decoupled from extensive host transcriptional reprogramming (Figure 5C) and cannot be solely explained by enhanced *Sv* growth (Figure 2F) as speculated for other fungal-bacterial synergistic beneficial effects (Del Barrio-Duque et al., 2019). Our transcriptional and phenotypic data further suggest that *Sv* – bacterial synergism in protecting host roots have also a component which is additive because the underlying antagonistic mechanisms displayed by the fungal root endophyte and the bacterial strains are likely to be distinct and explained mainly by direct microbe-microbe interactions outside the plant. Nonetheless we have observed a higher level of inter-kingdom mediated antagonism on *Bs* in presence of the host. This suggests a minor but relevant host-dependent effect which needs to be addressed (Figures 2A, 2E and 2F). At the early time point of 6 dpi, growth promotion was only observed in the combined presence of *Sv* and certain bacterial strains with the strongest effect during co-inoculation with Root11 in barley and Root172 in Arabidopsis (Figures 2C and S2). Furthermore, growth promotion required living microbes, as co-inoculation with heat-inactivated bacteria did not increase the root fresh weight in barley. Commensal bacteria in the rhizosphere can trigger plant growth promotion and resistance to pathogen (Pieterse et al., 2014; Souza et al., 2015; Vlot et al., 2020). Among them, strains belonging to the genus *Bacillus* are often used as bioagents due to their function in eliciting ISR (induced systemic resistance) as well as growth promotion (Kloepper et al., 2004; Vlot et al., 2020). However, plant growth promoting bacteria (PGB) and Sebacinales mediated growth promotion are often reported during later stages of colonisation. The early host growth enhancement observed with *Sv* and the bacteria might thus confer a competitive advantage for plants in nature. It is striking that the growth promoting effect is not accompanied by an extensive host transcriptional response with only 13 barley DEG being specific to this condition (Figure 5C; Table S7). Interestingly, several of these genes display differential expression across barley accessions (analysed using Genevestigator) compared to the cultivar Golden Promise. It would therefore be informative to test growth outcomes of combined *Sv* and e.g. Root11 inoculation in different barley varieties/ecotypes. The resulting synergistic inter-kingdom benefits in plant protection against fungal disease and in plant physiology (Figures 2 and 3) are in line with studies of the Sebacinales fungus *S. indica* with single bacterial strains on tomato (Del Barrio-Duque et al., 2019; Kumar et al., 2012; Sarma et al., 2011), rice (Dabral et al., 2020), barley (Varma et al., 2012) and chickpea (Mansotra et al., 2015) and underlay the broad functional relevance for fungi of the order Sebacinales in plant health in multi-kingdom environments.

The deployment of microbiota as biocontrol agents for crop protection and enhancement is an ancient concept (Vessey, 2003) which is gaining increased relevance in modern agriculture (Finkel et al., 2017; Vannier et al., 2019). Plant protection and growth promotion properties conferred by microbial consortia have been found to be more resilient than use of single strains (Finkel et al., 2017). Moreover, Duran et al. 2018 showed that a complex SynCom consisting of bacteria, fungi and Oomycetes led to strongest beneficial effects on Arabidopsis growth and survival compared to mono-kingdom or small SynCom associations and hypothesized that selective pressures over evolutionary time favor inter-kingdom microbe-microbe interactions over interactions with single microbial strains (Duran et al., 2018). Inter-kingom associations are frequently observed between members of the Sebacinales and bacteria. Different Sebacinales species host endobacteria of the orders *Bacillales* (genera *Paenibacillus*), *Pseudomonadales* (*Acinetobacter*) and *Actinomycetales* (*Rhodocuccus*) and its close relative *S. indica* hosts an endobacteria of the order *Rhizobiales* (*Rhizobium radibacter*) (Sharma et al., 2008). Beneficial effects of these intimate inter-kingdom interactions on the plant host and the fungus itself were described between *S. indica* and *R. radibacter* (Glaeser et al., 2016; Sharma et al., 2008) and for interactions between arbuscular mycorrhizal fungi and bacteria belonging to different species of the orders Proteobacteria (*Rhizobiales*) and Firmicutes (*Bacillales*) (Artursson et al., 2006). Considering the pervasiveness of beneficial effects conferred by *Sebacinales* and bacteria compared to the vulnerability of *Bs* in a multipartite context, our data support the hypothesis that establishment of beneficial inter-kingdom interactions in the plant microbiota is an evolutionary conserved and robust trait.

## Acknowledgments

We thank Paul Schulze-Lefert and the DECRyPT community (SPP 2125) for providing the bacterial strains used in this study. LM was supported by the Max-Planck-Gesellschaft through the International Max Planck Research School (IMPRS) on ‘Understanding Complex Plant Traits using Computational and Evolutionary Approaches’ and the University of Cologne. AZ and JEP acknowledge support from the Cluster of Excellence on Plant Sciences (CEPLAS) funded by the Deutsche Forschungsgemeinschaft (DFG, German Research Foundation) under Germany’s Excellence Strategy – EXC 2048/1 – Project ID: 390686111 and projects ZU 263/11-1 and PA 917/8-1 (SPP DECRyPT). JEP and CU acknowledge The Max Planck Society for additional support. Graphical illustrations were designed with the BioRender online tool. SM would like to express our gratitude to Prof. Francis Martin for allowing us to use the secretome prediction and visual omics workflow on the computing cluster at INRAE Nancy, France.

## Supplementary Figures

**Fig. S1:**
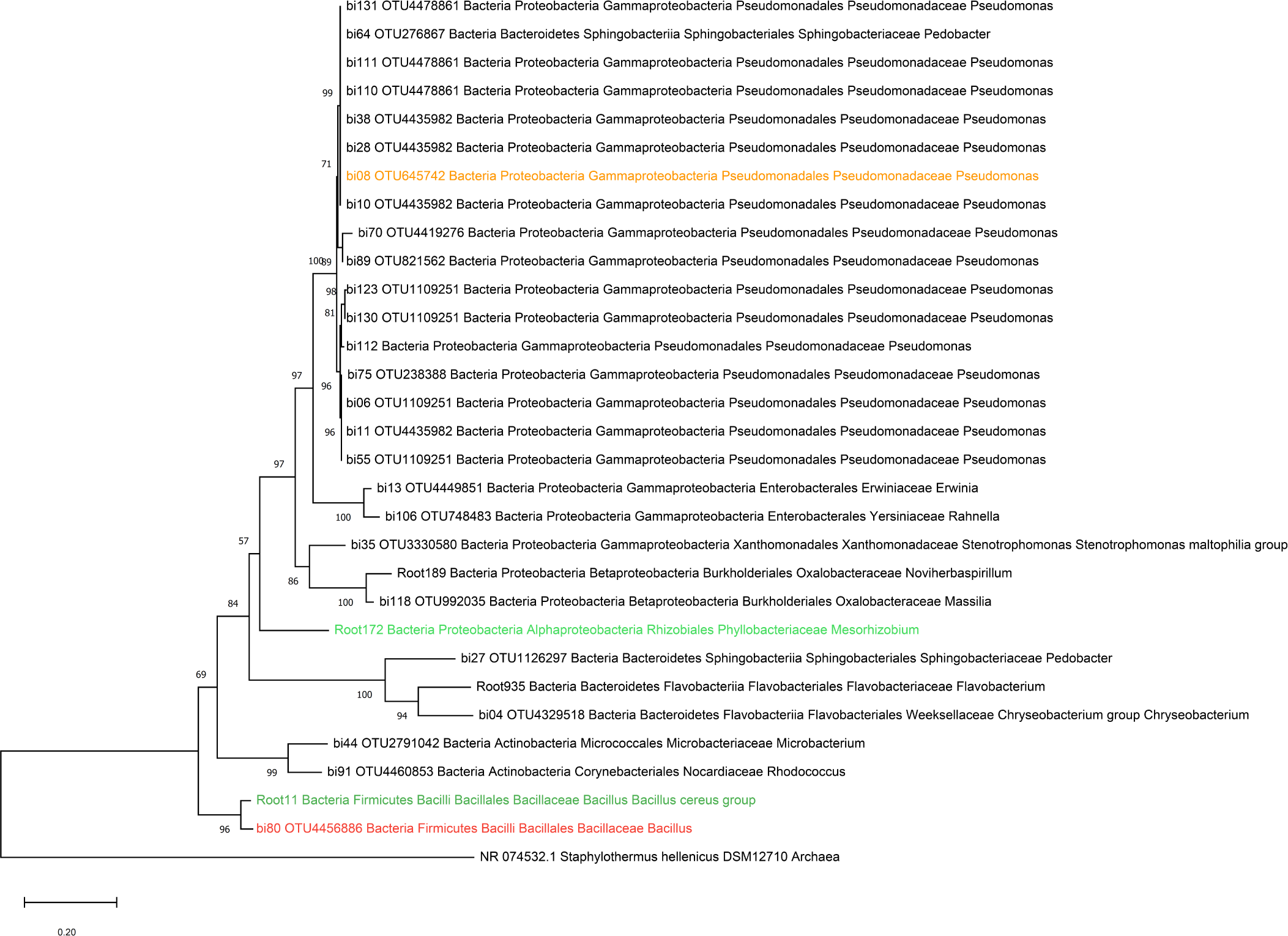
Phylogenetic tree of Arabidopsis and barley associated bacteria. The evolutionary history was inferred from 16S rRNA genes (Bai et al., 2015) by using the Maximum Likelihood method and Tamura-Nei model (Tamura et al., 1993). The percentage of trees in which the associated taxa clustered together is shown next to the branches. The tree is drawn to scale, with branch lengths measured in the number of substitutions per site. Evolutionary analyses were conducted in MEGA X (Kumar et al., 2018). Taxonomy of strains was inferred by blast searches against NCBI rRNA/ITS databases.

**Fig. S2:**
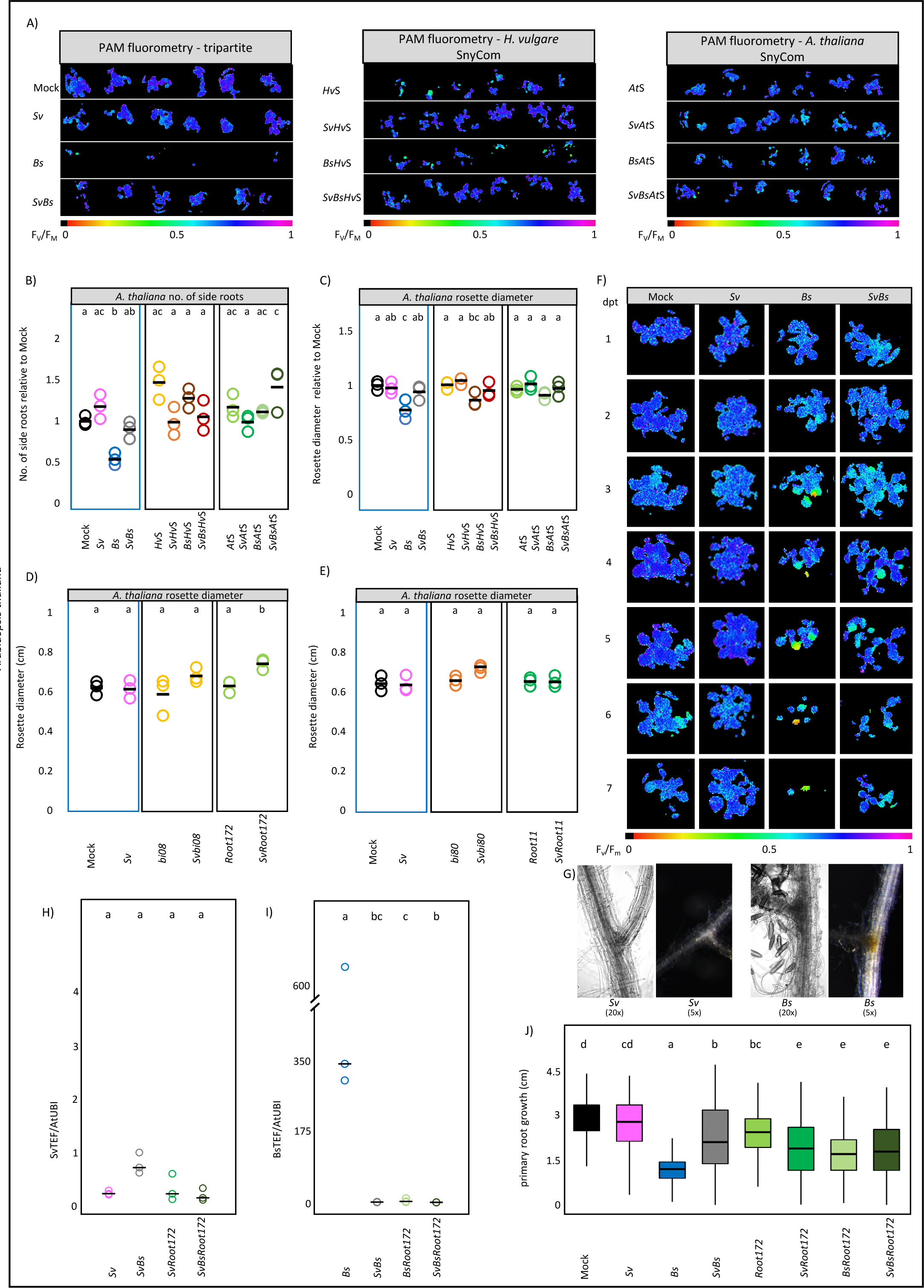
Phenotypic analysis of Arabidopsis roots at 6 dpi with *Sv* and/or *Bs* with or without the bacterial SynComs *Hv*S and *At*S or single bacterial strains. A) The photosystem II (PSII) quantum yield of 5 At seedlings/well in absence or presence of *Sv*, *Bs* and/or a bacterial SynCom (*Hv*S or *At*S) at 4 dpt after dark adaptation (F_V_/F_M_) via PAM fluorometry. Purple/dark blue, lighter colors and black color indicate high, reduced and lack of PS II activity respectively. B) number of *A. thaliana* side roots relative to control plants (Mock) C-E) *A. thaliana* rosette diameter in presence or absence of *Sv*, *Bs* and the different bacterial strains/SynComs and C) the bacterial SynComs. D) the single Proteobacteria strains bi08 and Root172 E) the single Firmicutes strains bi80 and Root11. F) exemplary time cause PAM fluorometry pictures from 1-7 d post transfer in the tripartite conditions G) Pictures of *Sv* and *Bs* inoculated Arabidopsis roots at 6 dpi in 5x and 20x magnification H) *Bs* and H) *Sv* colonisation in Arabidopsis inoculated with *Sv*, *Bs* or both fungi in the absence of presence of Root172 at 6 d post inoculation inferred by expression analysis of the fungal housekeeping gene TEF compared with Arabidopsis ubiquitin (UBI) (n = 3 with 60 plants per replicate). I) Root growth of Arabidopsis seedlings inoculated with Sv, Bs or Root172 in all combinations from 0 dpi – 6 dpi (cm). Different letters represent statistically significant differences according to one-way ANOVA and Tukey‘s post-hoc test (p < 0.05).

**Fig. S3:**
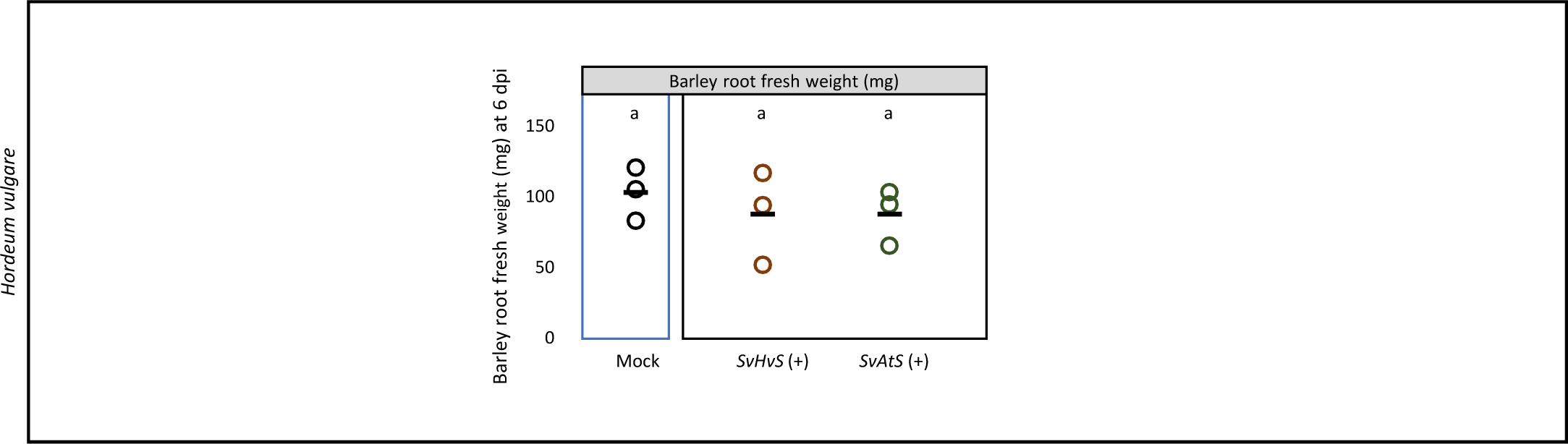
Root fresh weight (mg) of barley seedlings inoculated with *Sv* and the heat-inactivated bacterial SynComs (+). Root weight was measured at 6 dpi. Different letters represent statistically significant differences according to one-way ANOVA and Turkey‘s post-hoc test (p < 0.05).

**Fig. S4:**
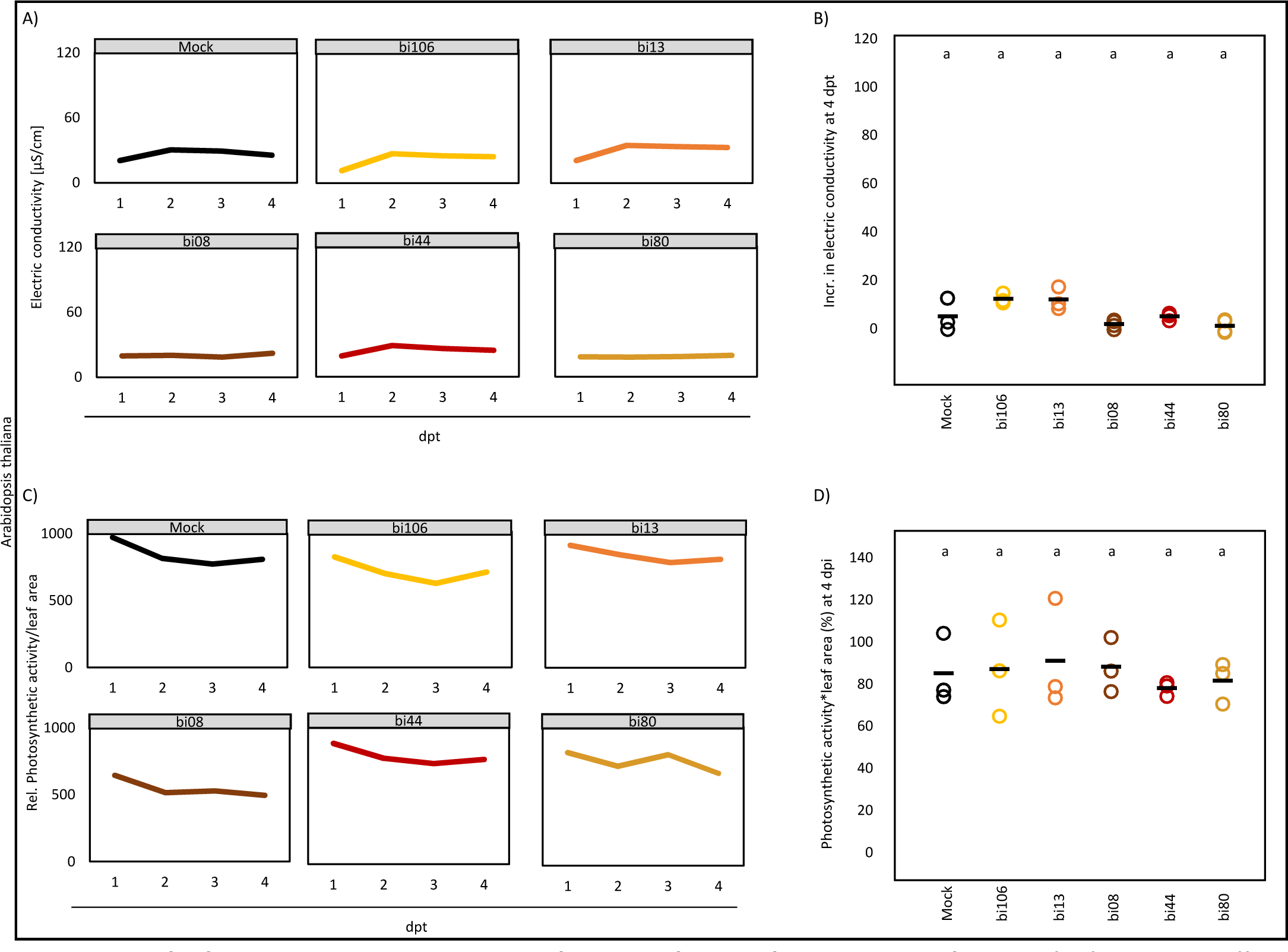
Arabidopsis root responses to bacterial inoculations at 6 dpi. Arabidopsis seedlings were inoculated with individual strains bi106, bi13, bi08, bi44 and bi80, all derived from the *Hv*S. A) Electric conductivity from 1 to 4 days post transfer (n = 6). B) Total increase in electric conductivity from 1 to 4 days post transfer (n = 3). C) Photosynthetic activity (F_V_/F_M_) from 1 to 4 days post transfer (n = 3). D) Photosynthetic activity per leaf area at 4 dpi relative to 1 dpi (n = 3). Different letters represent statistically significant differences according to one-way ANOVA and Tukey‘s post-hoc test (p < 0.05).

**Fig. S5:**
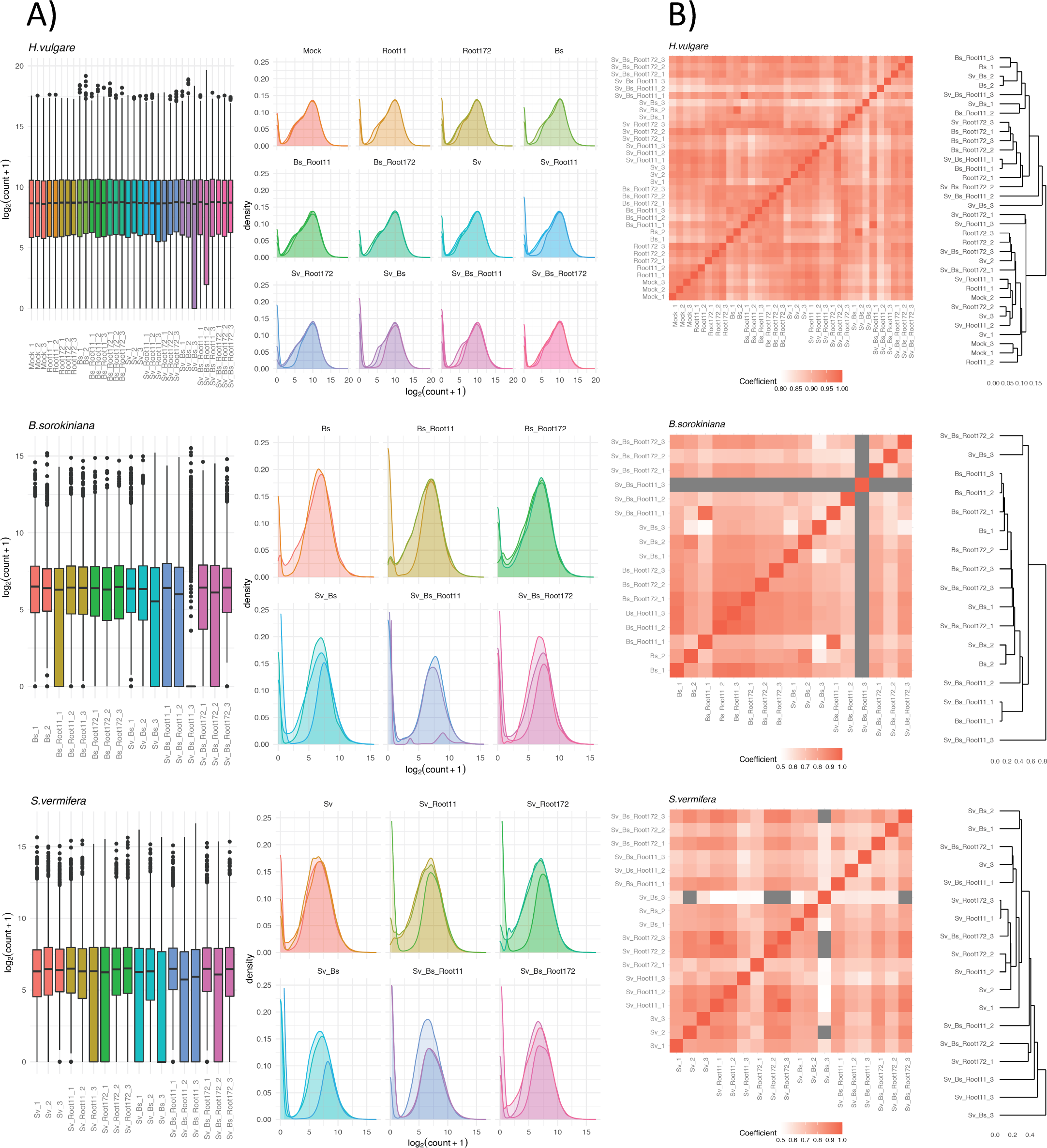
Assessment of RNA-seq data. A) Distribution and density of the normalised log2 transformed transcript count of genes for three organisms. B) Correlation of transcriptomes of RNA-seq samples. Left part: Adjacent matrix based on the of the correlation coefficients. Right part: Hierarchical clustering of biological replicates according to the distances of transcriptomic similarities. Grey represents correlation coefficients lower than 0.5.

**Fig. S6:**
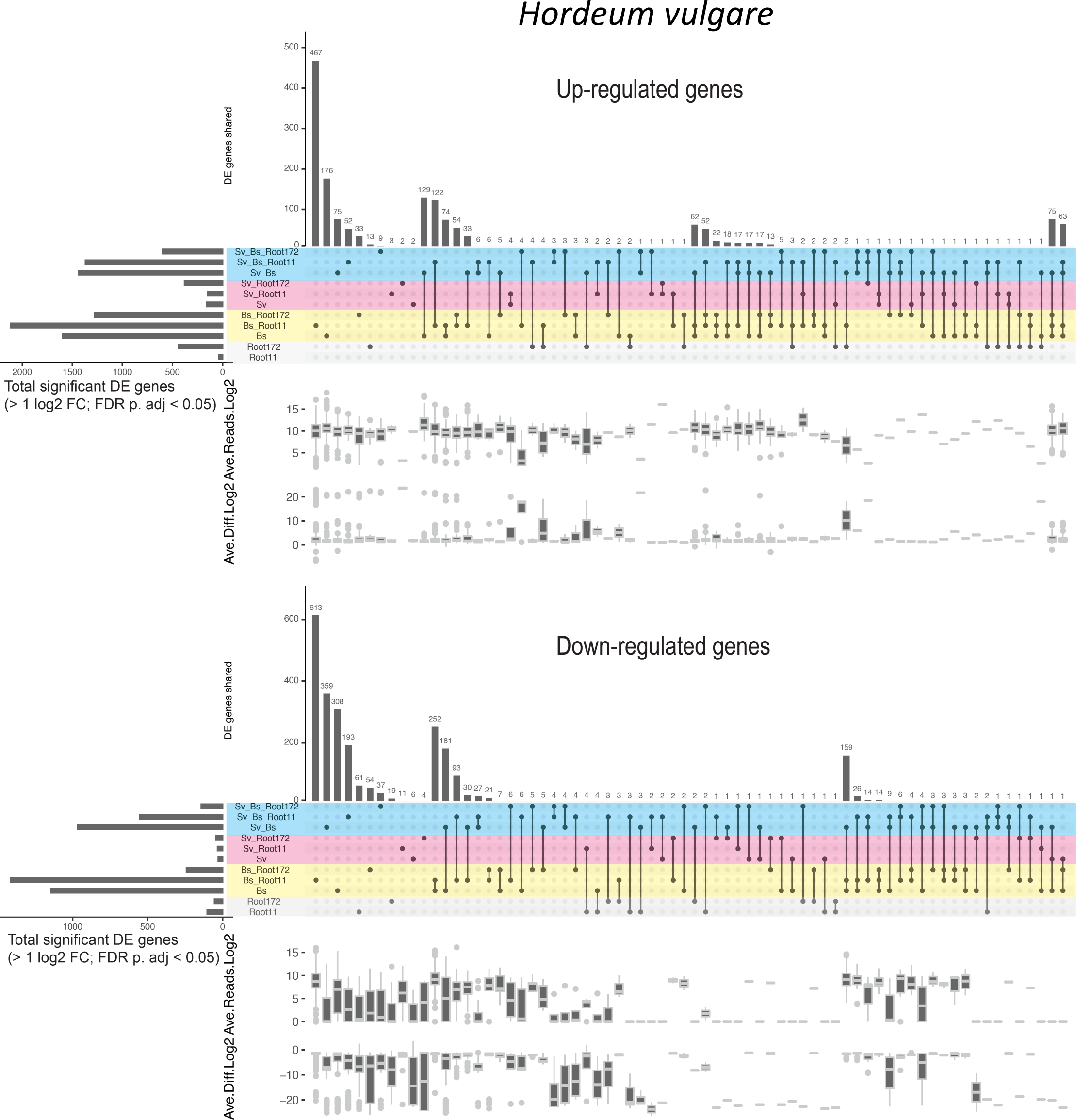
*H. vulgare* differentially expressed genes. Selected DEGs (> 1 log2FC; FDR adjusted p-value < 0.05) are compared to barley Mock control. Up and down-regulated genes are separately presented (see the combined figure, main Fig. 5C). Horizontal bars: Total number of DEGs per condition. Vertical bars: Number of genes unique/shared for top 70 intersections. See Table S7.

**Fig. S7:**
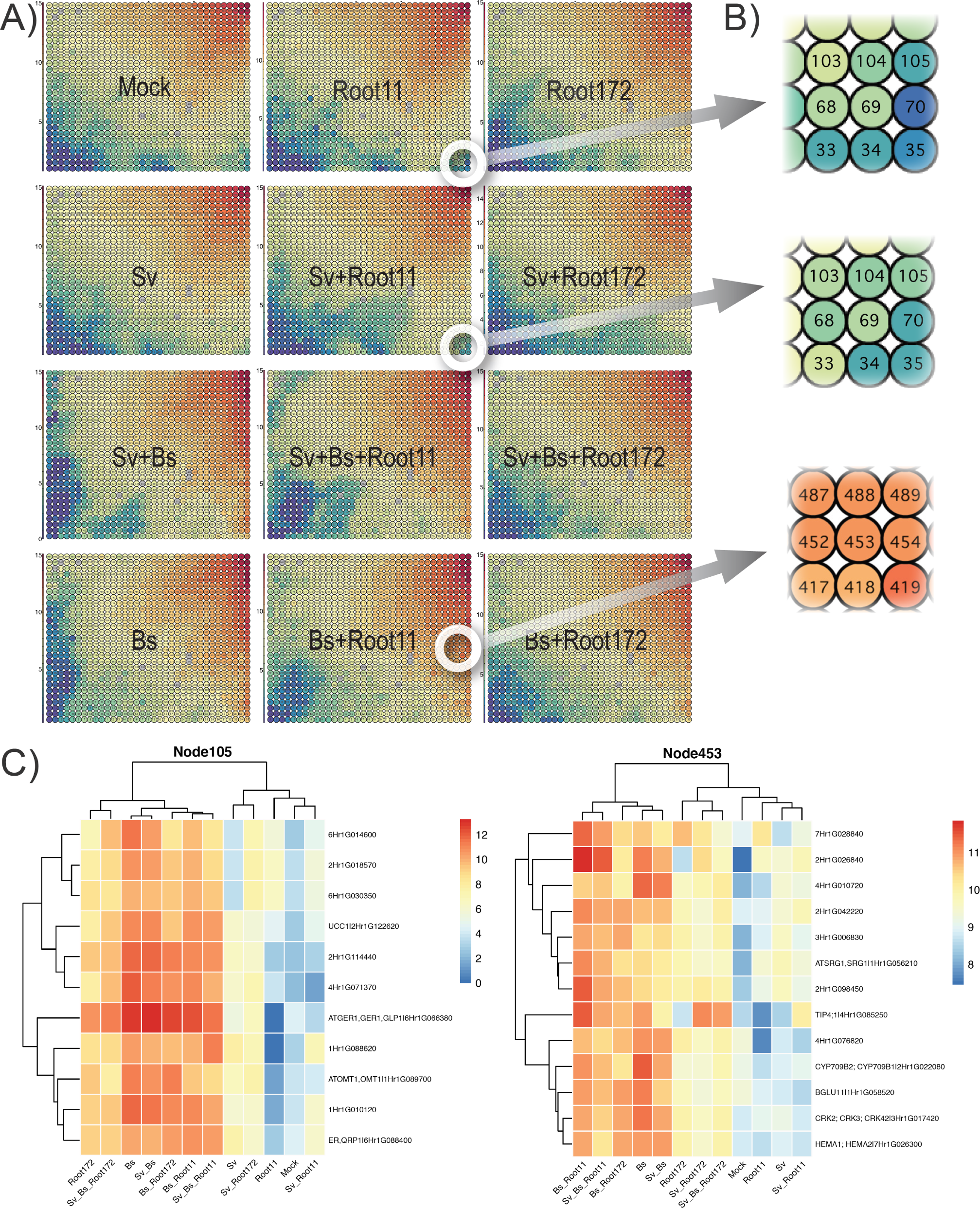
Genome-wide transcriptomic dynamics of *H. vulgare* per condition. A) Trained Self-Organizing Maps (SOM, Tatami maps) showing barley global transcriptomic trends. Colors indicate the averaged log2 read count of replicates from each of the conditions. Each circle represents a node (IDs 1 to 1015). Single nodes contain approximately 10 to 100 genes. The SOM resulted in similarly-expressed genes separated into high, medium, and low expressed groups. The highly transcribed genes are clustered at the top right corner (red) and the lowly transcribed groups at the bottom left corner (blue). Barley inoculated with *S. vermifera* (*Sv*) exhibited similar patterns to barley mock. The presence of the pathogen (*Bs*) was a major factor driving responses in the host, which was consistent with the dynamics of DEG shown in Fig. 5. There were additional effects of the co-inoculated bacteria on barley (Root11 and Root172). The shape of the lowly transcribed clusters shifted in co-inoculated roots with the bacterial strains (e.g. *Bs* vs *Bs*+Root11 or *Bs*+Root172). B) Double-circles (i.e. white doughnuts) on Tatami maps indicate the location of highly regulated gene groups and such gene groups are magnified. C) Examples of highly regulated genes (FDR adjusted p < 0.05) present in particular nodes. The high and low log2 gene expression is displayed in red and blue respectively. Gene identification number with corresponding annotations (if there is any) are presented on Y-axis. Node 105 contains similarly lowly expressed genes for barley mock, bacterium 11, and *S. vermifera* (Mock, Root11, Sv_Root11). Node 453 shows highly expressed genes for *B. sorokiniana* with bacterium 11 (Bs_Root11). See Table S3 for details.

**Fig. S8:**
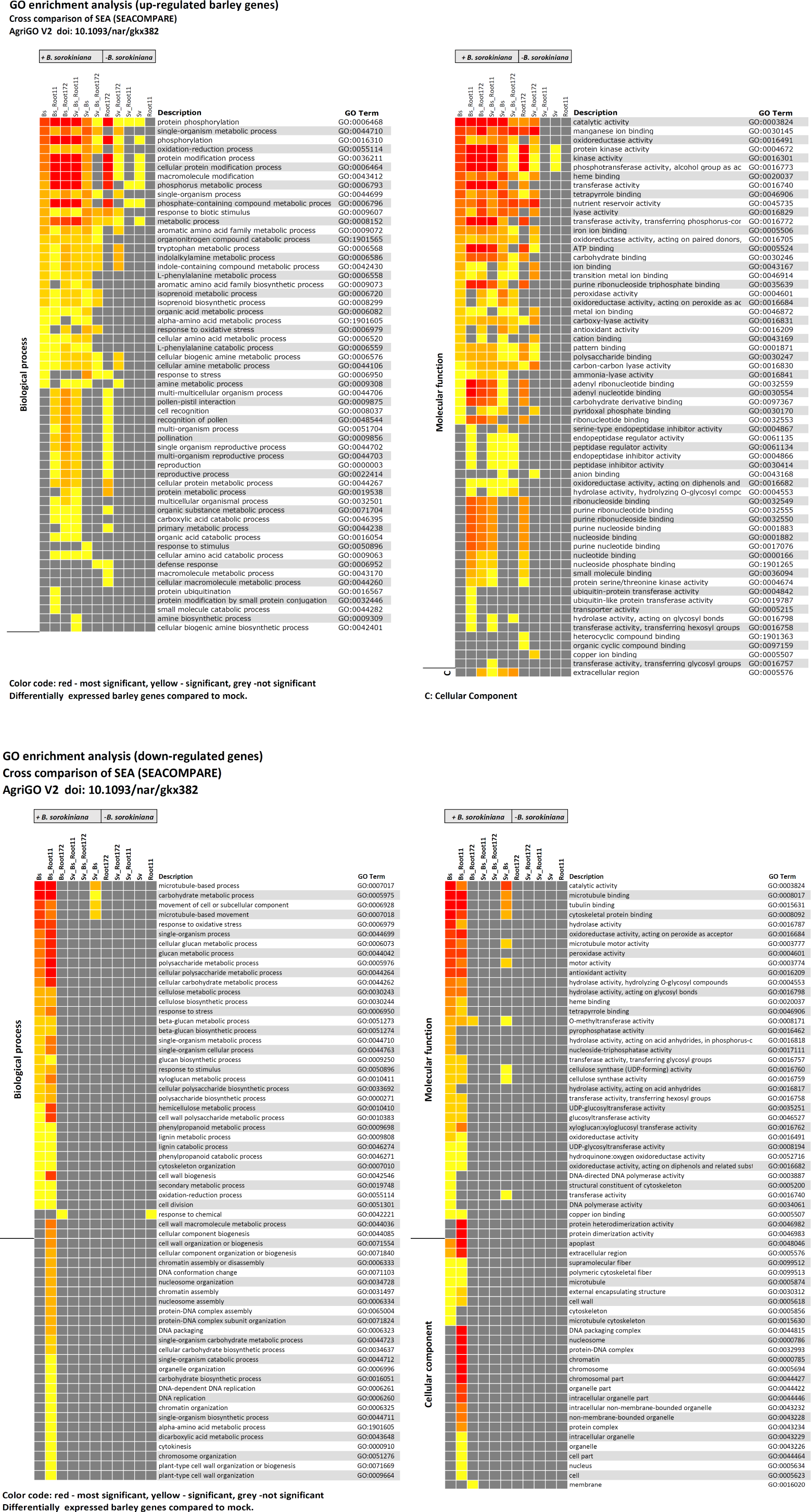
GO enrichment analysis (up- and down-regulated barley genes)

**Fig. S9:**
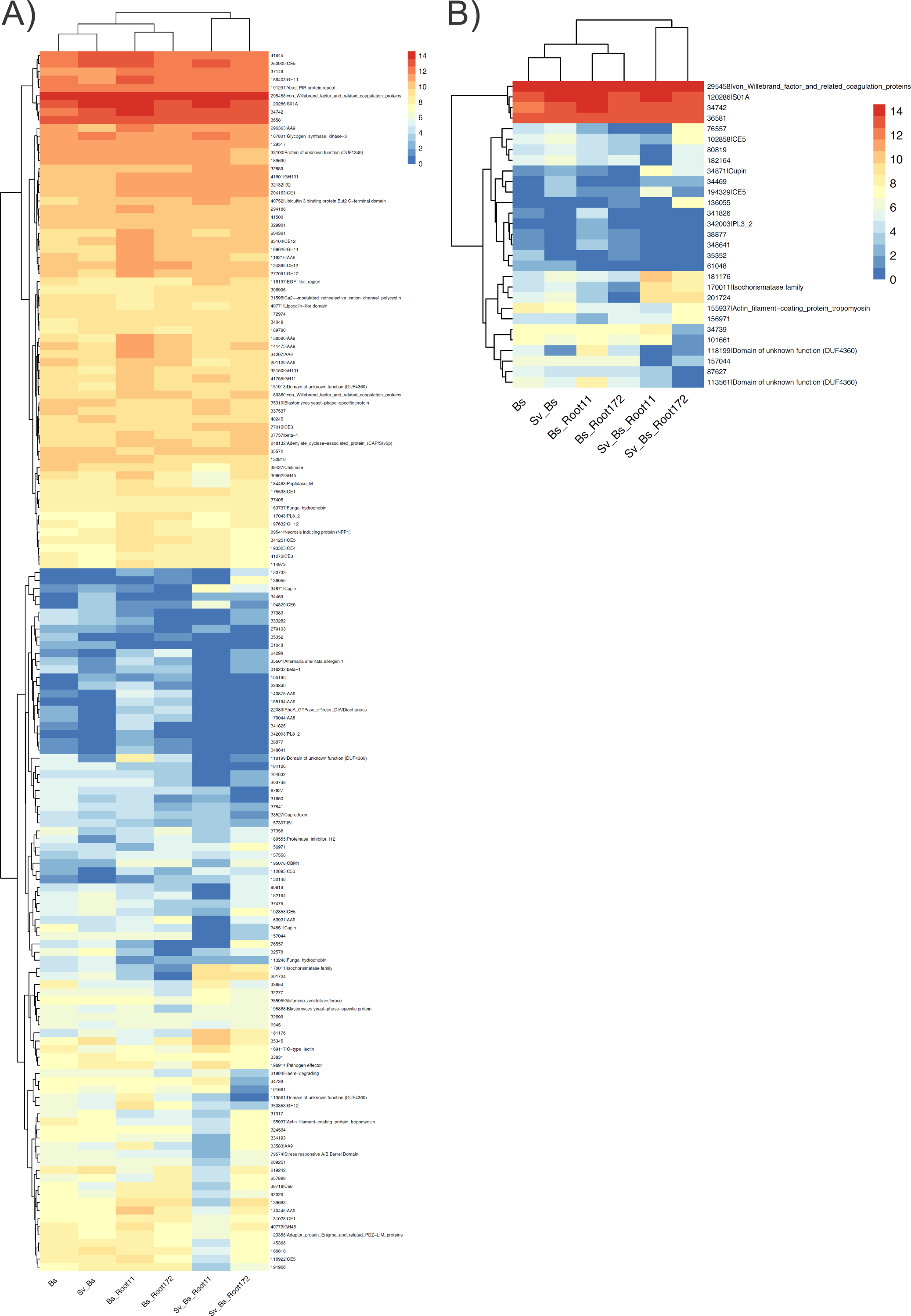
Expression of genes coding for effectors in *B. sorokiniana*. A) Averaged log2 read count of genes under the conditions. Y-axis shows JGI Protein IDs with corresponding annotations. B) Averaged log2 read count of genes with high loadings (see Methods). Y-axis shows JGI Protein IDs with corresponding annotations if there is any. See Table S9.

**Fig. S10:**
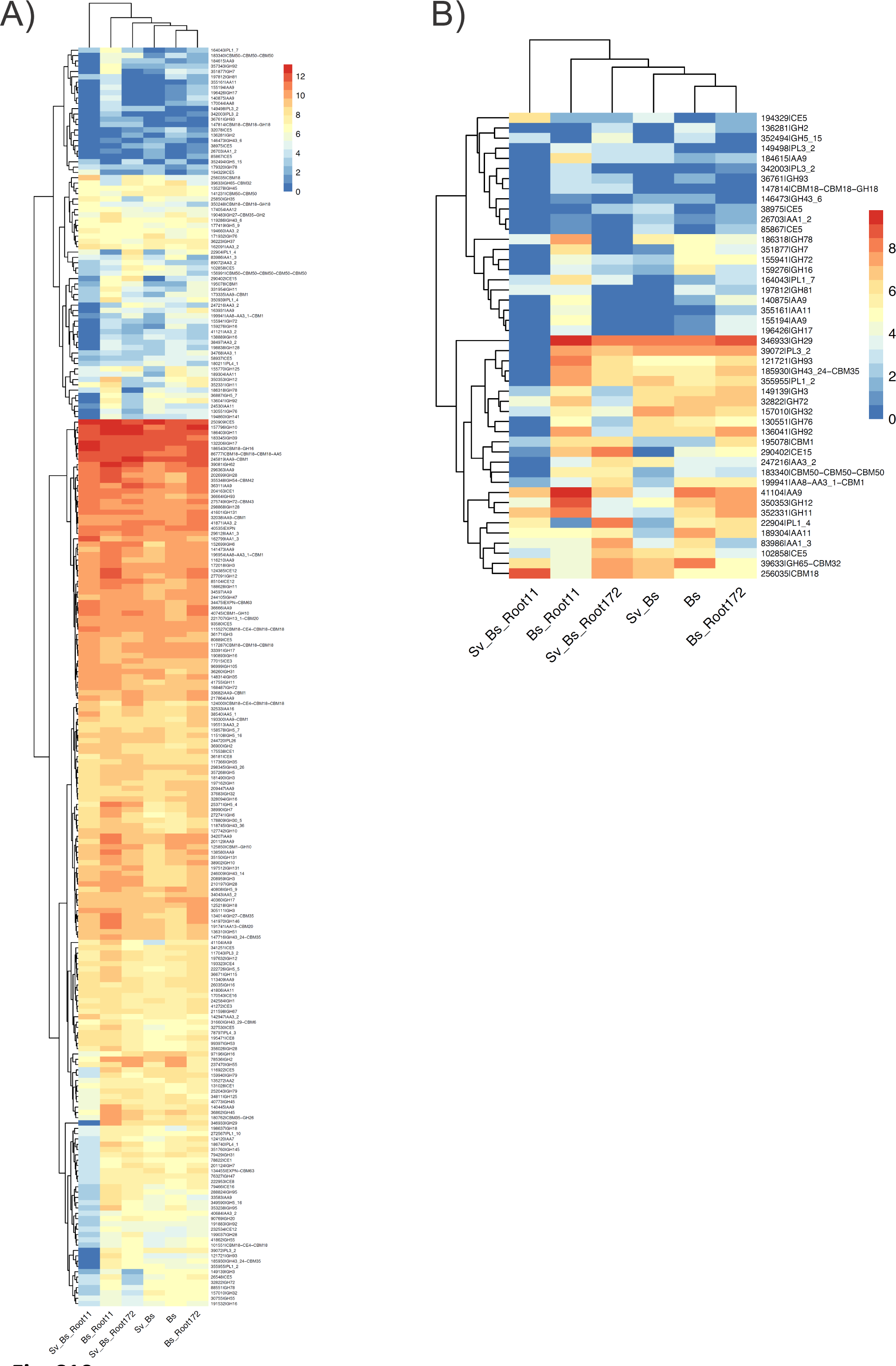
Expression of genes coding for CAZymes predicted to be secreted in B. *sorokiniana*. A) Averaged log2 read count of genes under the conditions. Y-axis shows JGI Protein IDs with corresponding annotations. B) Averaged log2 read count of genes with high loadings (see Methods). Y-axis shows JGI Protein IDs with corresponding annotations if there is any. See Table S9.

**Fig. S11.**
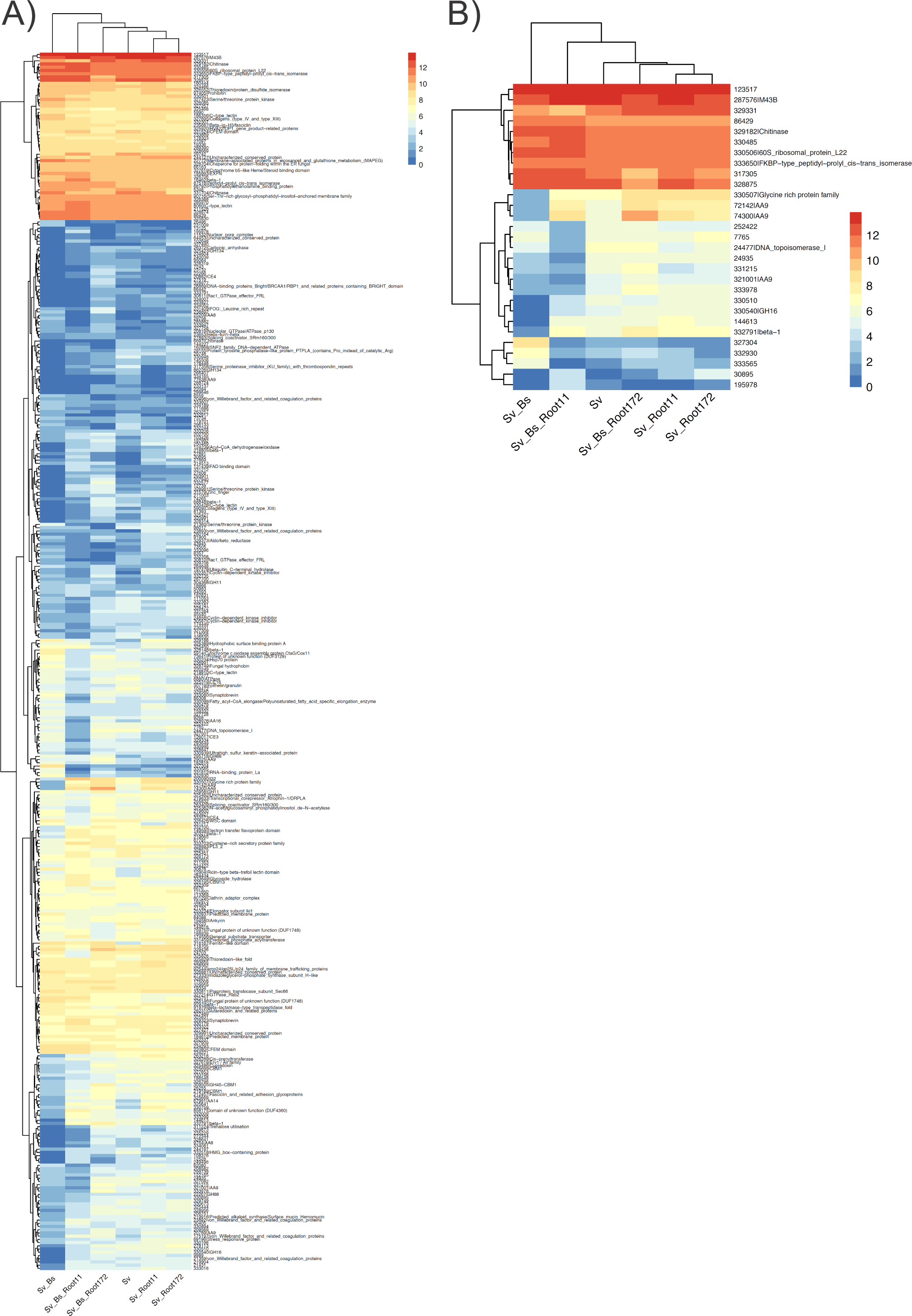
Expression of genes coding for effectors in *S. vermifera*. A) Averaged log2 read count of genes under the conditions. Y-axis shows JGI Protein IDs with corresponding annotations. B) Averaged log2 read count of genes with high loadings (see Methods). Y-axis shows JGI Protein IDs with corresponding annotations if there is any. See Table S9.

**Fig. S12:**
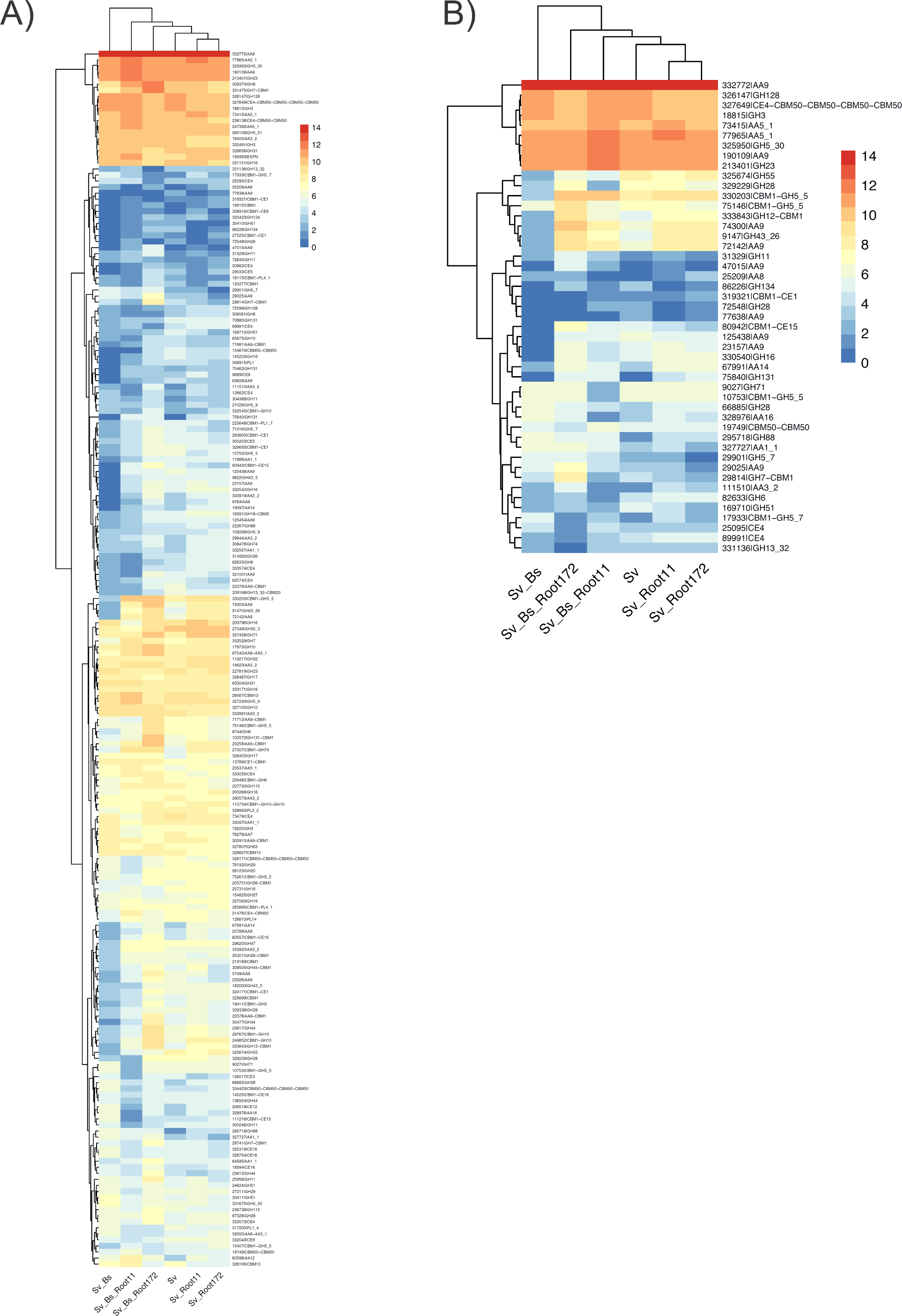
Expression of genes coding for CAZymes predicted to be secreted in *S. vermifera*. A) Averaged log2 read count of genes under the conditions. Y-axis shows JGI Protein IDs with corresponding annotations. B) Averaged log2 read count of genes with high loadings (see Methods). Y-axis shows JGI Protein IDs with corresponding annotations if there is any. See Table S9.

**Fig. S13:**
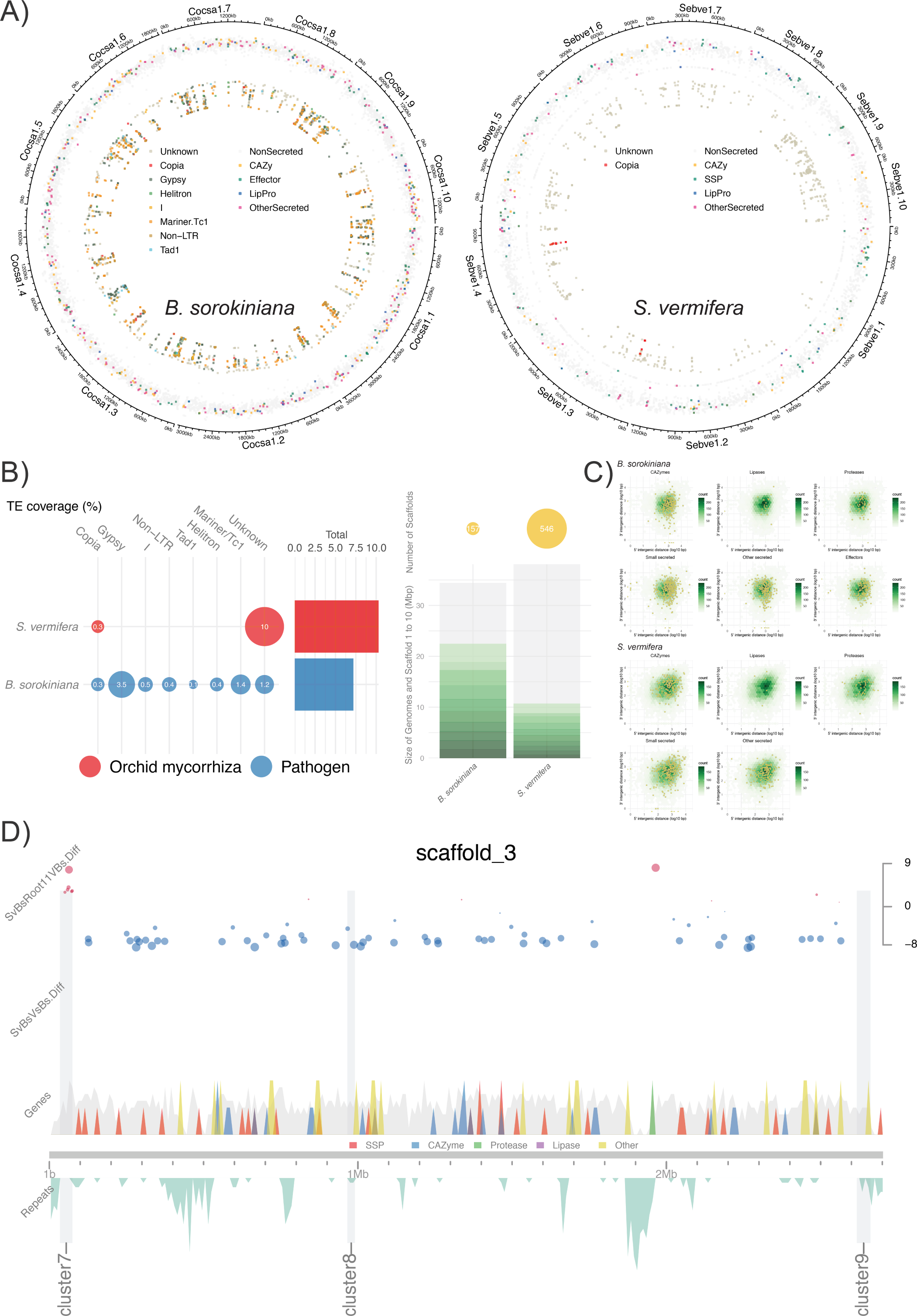
Genomic features of *B. sorokiniana* and *S. vermifera*. A) The genomic location of genes and transposable elements (TEs) are visualised with the largest 1 to 10 scaffolds from the genome assemblies. Hanabi plots (fireworks in Japanese) contains three rings. Outer ring: The size of scaffold 1 to 10 presented clock-wise starting from 3 o’clock. Colors of Scaffold 1 to 10 are from dark grey to light grey. The boxes next to “fungal names + scaffold ID” represents the length of the scaffolds. Approximate locations of genomic features can be seen with the small rulers aligned in the outer ring. Middle ring: The genomic locations of all genes based on JGI GFF files. Genes coding for theoretically secreted proteins (CAZymes, SSPs, lipases, proteases) are in color. Other genes coding for non-secreted (i.e. intracellular) proteins are in grey. Inner ring: The genomic locations of TE families and unidentified repeats. Repeat sequences (>50 bases with >10 occurrences in a genome) were identified. Vertical axis for the density of genes/TEs in the rings: The mean distance of neighboring genes or TEs in log2. If distances between genes/TEs are short, dots (i.e. the locations of genes and TEs) go towards the centre of plots. If distances between genes/TEs are long, dots go towards the outer circle (it gives a sense of how densely localized or dispersed genes/TEs are). See Table S10 for details. B) TE content and scaffolds in the genome assemblies. Left panel: Coverage of transposable elements in the genomes. The size of the bubbles corresponds to the percentage of TE coverage in the genomes. Right panel: Genome size and the number of scaffolds. The bars in grey indicate the genome size. Individual green sections shows the largest scaffolds 1 to 10. The circle size corresponds to the number of total scaffolds. The ecological lifestyle is in color. C) Intergenic distances of genes for secreted proteins (i.e. intergenic distance = gene to gene distance). Proteins predicted to be secreted are categorised into CAZymes, proteases, lipases, the rest of secreted protein, effectors, and a subcategory for small secreted proteins (< 300 amino acids). Yellow points: Intergenic 5’ and 3’ distances of individual genes. Green tiles: Density of intergenic distances of all genes present in a genome. Genes tend to be gathered at the centre of the maps, showing average intergenic distances. Genes nearby a cluster of transposable elements tend to show long intergenic distances (see top right corner) where new functions of genes might be evolved due to the transposition. See Table S11. D) Visual integration of multi-omics showing highly regulated biosynthetic gene clusters in *B. sorokiniana*. Omics data (transcriptome, secretome, repeatome and genome) are combined and visualised. Scaffold 3 from the genome assembly is presented for example. Grey vertical bars: Biosynthetic gene clusters. Top panel: Significantly regulated genes under conditions. The size of circles and colors correspond to differential transcription levels in log2. Middle panel: The genomic locations and density of all genes (grey) and gene for secreted proteins (colors). The scaffold size of a genome assembly is shown as a grey horizontal bar. Bottom panel: The genomic location and density of total and individual TE families. See Table S12 for details.

## Supplementary Methods to Fig. S13

*Multi-omics integration and visualization for fungi.* Secreted proteins were predicted using the method described previously (Pellegrin *et al*., 2015). CAZy annotations were provided from CAZy team (www.cazy.org). Transposable element (TE) identification was performed with Transposon Identification Nominative Genome Overview (TINGO; Morin *et al*., 2019). We predicted biosynthetic gene clusters with antiSMASH 5.1 (Madema *et al*., 2011). Differential expression of genes was calculated with the control, *B. sorokiniana* alone grown in barley using DESeq2 (Love *et al*., 2014). We excluded genes showing either very low raw reads or adjusted p value (FDR) larger than 0.05. Differentially expressed genes coding for effectors were obtained from the previous study (Sarkar *et al*., 2019). Output files obtained from the various analyses above and functional annotations from JGI MycoCosm were cleaned, sorted, combined and visualized using a set of custom R scripts, Visually Integrated Numerous Genres of Omics (VINGO; Looney *et al*., 2021) incorporating R package karyoploteR (Gel & Serra 2017). Also, we located genomic features (i.e. genes, predicted secretome, transposable elements) in the largest scaffold 1 to 10 in a circular manner (Hanabi plots) with Syntenic Governance Overview (SynGO; Hage *et al*., 2021) incorporating R package Circlize for visualization (Gu *et al*., 2014).

*Visual intergenic distances in genomes with statistics.* Intergenic distances in the genomes were calculated based on the study (Saunders *et al*. 2014). The original scripts are obtained from https://github.com/Adamtaranto/density-Mapr. Theoretically secreted proteins were determined with Secretome pipeline mentioned above. The results were visualized using a visual pipeline SynGO (Hage *et al*., 2021). The mean TE-gene distances were calculated from; (i) the locations of observed genes and TEs; and (ii) random “null hypothesis” genome models made by randomly reshuffling the locations of genes. The distribution of genomic features was purely random for null models and there was no association between the locations of genes and repeat elements. The probability (p-value) of mean TE-gene distances was calculated based on a normal distribution of 10,000 null hypothesis models. The process was performed with R package, regioneR (Gel *et al*., 2016).

